# A premitotic polarity program patterns the grass leaf epidermis

**DOI:** 10.64898/2026.07.07.736711

**Authors:** Anastasiia L. Korosteleva, Kim Janssen, Dan Zhang, Paola Ruiz Duarte, Irem Polat, Nika Goršek, Barbara Jesenofsky, Elif Baç, Heike Lindner, Michael T. Raissig

## Abstract

Transverse asymmetric cell divisions (ACDs) in grass leaf epidermal development produce large basal pavement cells and small apical specialised cells. These “patterning divisions” generate the long-short epidermal cell pattern that is distinctive of grasses. Here, we show that patterning divisions require premitotic basal polarisation of Bd-POLAR-LIKE1 (BdPL1) in the model grass *Brachypodium distachyon*. Loss of *BdPL1* disrupted division-plane orientation and postmitotic cell-size asymmetry in all cell files, which resulted in epidermal patterning defects. Ectopic expression analyses demonstrated that BdPL1 polarisation was independent of cellular context and sufficient to promote supernumerary transverse divisions. Furthermore, the developmental regulators BdBREVIS RADIX-solo and BdYODA1 formed a post-division polarity domain enforcing cell fate asymmetry independently of BdPL1. We propose that the premitotic BdPL1 module enforces physical cell-division asymmetry contributing to medio-lateral patterning of cell types, whereas the postmitotic BdBRX-solo/ BdYDA1 module enforces within-file cell fate asymmetry. Together, they robustly pattern the grass leaf epidermis.

## Introduction

Cell polarity is a fundamental biological principle that enables asymmetric cell divisions (ACDs) and facilitates the distinct postmitotic identities of the daughter cells. Particularly in plants, where complex organ development occurs in the absence of cell migration, accurate temporal and spatial control of cell division patterns is essential for organ and tissue development^1–3^. Extrinsic, axial and intrinsic polarity cues position the division plane and establish unequal distribution of cell fate determinants to enforce both cell size (i.e., physical) and cell fate asymmetry^3,4^. Although major progress has been made in identifying general and lineage-specific polarity factors in *Arabidopsis thaliana*, with extensive research on the stomatal lineage^2,4–6^, less is known about how polarity mechanisms have diversified in grasses, particularly in the context of the distinct pattern and ontogeny of the grass leaf epidermis.

In contrast to broad-leafed eudicot plants like *A. thaliana*, where the planar leaf surface grows both longitudinally and medio-laterally, the grass leaf grows predominantly along the longitudinal axis, which is reflected in the cellular pattern of its epidermis. The grass leaf epidermis is organised in vertical rows of semi-clonal cells, also referred to as cell files, that follow an acropetal developmental gradient from undifferentiated protodermal cells at the base of the leaf to mature specialised cells at the top (Figure 1A)^7–9^. Furthermore, developing epidermal cells divide only in two perpendicular orientations in grasses, transverse and longitudinal. In the wild model grass *Brachypodium distachyon*, stage 1 protodermal cells divide symmetrically, and cell files are medio-laterally patterned into stomatal or hair cell files^8,10^. At this stage, stomatal cell file identity or hair cell identity is antagonistically established by *SPEECHLESS1* (*SPCH1*), *SPCH2* and *INDUCER of CBF EXPRESSION1* (*ICE1*), or by *SQUAMOSA-PROMOTER BINDING PROTEIN-LIKE* (*SPL*) proteins, respectively^10–12^. In stage 2, transverse asymmetric divisions generate a smaller apical cell and a larger basal cell^7,9^. This division is also termed “patterning division” as it establishes the stereotypical long-short pattern of grass epidermal cell files, where a small specialised cell (e.g. stomata or hair cell) is interspersed by at least one large pavement cell^8,13^. Small apical cells obtain a specialised cell fate depending on the previously determined cell file identity and form guard mother cells (GMCs) or hair cell precursors in stage 2^10–12,14^. Hair cell precursors immediately grow perpendicular to the leaf surface using a tip-growth program^15^. In stomatal cell files, however, GMCs express the transcription factor MUTE, which moves into lateral neighbour cells to establish subsidiary mother cell (SMC) identity in stage 3^16–18^. SMCs then polarise towards the GMC, before a longitudinal ACD produces a lateral subsidiary cell (SC) during stage 4^19,20^. Finally, at stage 5, the GMC divides symmetrically to form a pair of guard cells (GCs), and the late-acting FAMA and SCREAM2 (SCRM2) transcription factors guide pore formation, differentiation and morphogenesis of the dumbbell-shaped grass GCs^10,12,21–23^.

**Figure 1.**
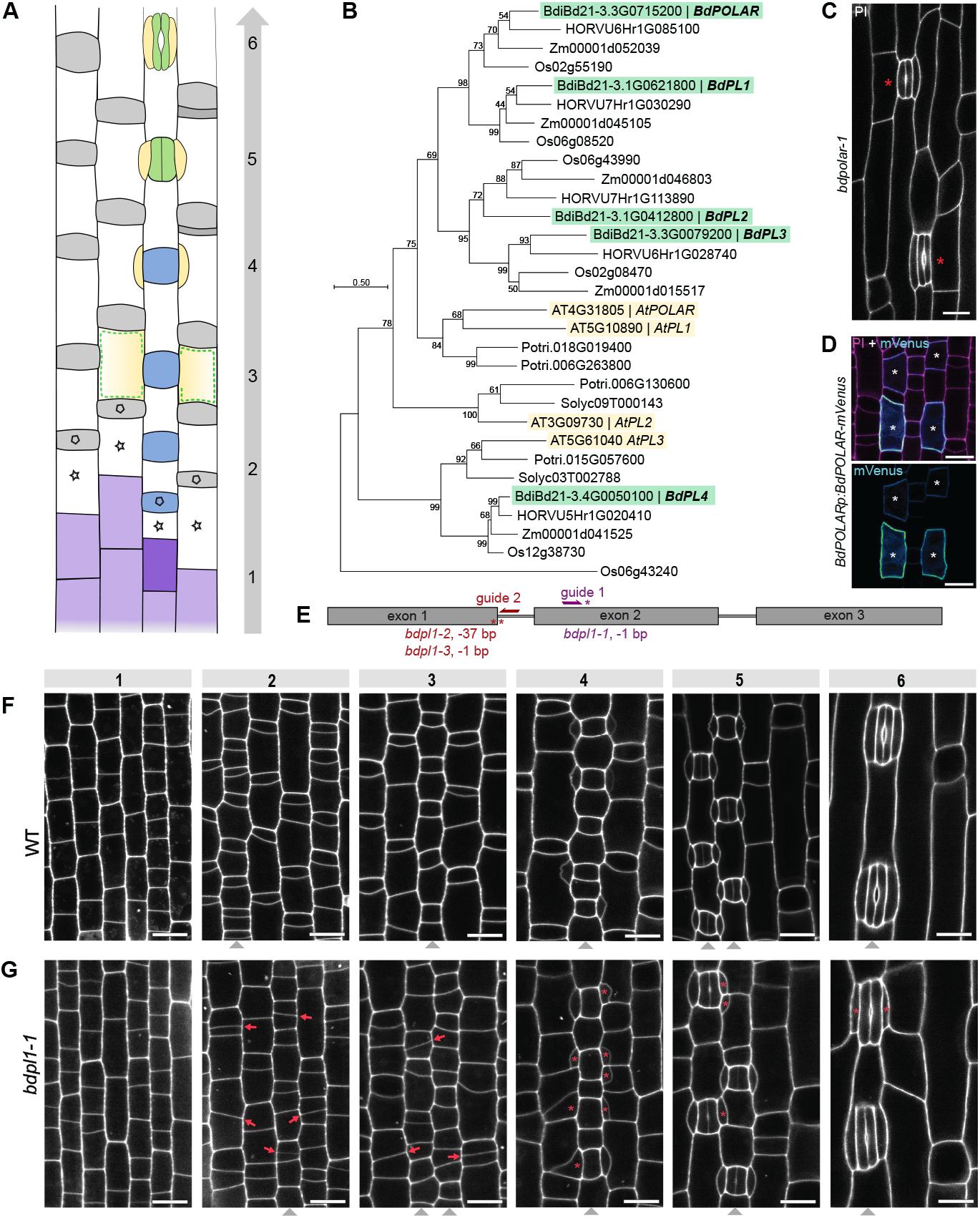
*BdPL1* controls the transverse asymmetric patterning division. **(A)** Leaf development in *Brachypodium distachyon*. (1) Transverse, symmetric divisions of protodermal cells, where stomatal (dark purple) and non-stomatal (light purple) cell files are defined; (2) transverse asymmetric cell divisions (ACDs), basal cells marked with a star, apical cells marked with a pentagon; (3) guard mother cell (blue) induces neighbouring cells to become subsidiary mother cells (SMCs), (4) longitudinal ACDs in SMCs produce subsidiary cells (yellow). Additional transverse ACDs in non-stomatal cell files above veins can produce silica cell precursors below hair cell (dark grey and light grey, respectively); (5) symmetrical, longitudinal division in GMCs forms two guard cells (green), (6) differentiation of the stomatal complex. **(B)** Phylogenetic tree of the *POLAR* family across seven species (*B. distachyon, O. sativa, H. vulgare, Z. mays, A. thaliana, S. lycopersicum* and *P. trichocarpa*). *B. distachyon* genes highlighted in green, *A. thaliana* genes highlighted in yellow. **(C)** Mutant phenotype of *bdpolar-1* with defective SC divisions (indicated with red asterisk)^33^. **(D)** *BdPOLARp:BdPOLAR-mVenus* expression in SMCs (marked with white asterisks)^33^. The upper panel shows merged signals of mVenus and propidium iodide (PI), lower panel shows only mVenus signal. **(E)** *BdPL1* gene model with the two CRISPR/Cas9 guides and induced mutations indicated. **(F-G)** Six developmental stages of the 2^nd^ leaf epidermis of 5-day-old wild-type (WT) (F) and *bdpl1-1* (G) seedlings. Stages are indicated according to the scheme in (A). Cell walls stained with propidium iodide (PI). Red arrows point at misoriented divisions, red asterisks mark defective SCs. Grey arrowheads indicate stomatal cell files. Scale bars, 10 μm.

In grasses, epidermal patterning divisions are spatially and temporarily restricted at the developmental zone at the base of the leaf^7,9^. In broad-leafed eudicots like *A. thaliana*, however, new divisions can occur throughout the leaf epidermis. In *A. thaliana*, stomatal lineage precursor cells (i.e., meristemoid mother cells) are established seemingly in a stochastic manner and can undergo between one and three ACDs before committing to GMC identity. Furthermore, the bigger stomatal lineage ground cell can either differentiate into a pavement cell or re-establish division capacity and re-iterate stomatal ACDs^24^. Epidermal ACDs in *A. thaliana* are governed by polarity proteins such as BREAKING OF ASYMMETRY IN THE STOMATAL LINEAGE (BASL)^25^, BREVIS RADIX (BRX) family proteins^26^, OCTOPUS-LIKE proteins (OPLs)^27^ and POLAR LOCALISATION DURING ASYMMETRIC DIVISION AND REDISTRIBUTION (POLAR) family proteins^28,29^. Several of these polarity proteins act as scaffolds for downstream signalling effectors, whose asymmetric distribution contributes to attenuating the activity of the ACD-promoting transcription factor AtSPCH. AtPOLAR, for example, scaffolds GSK3-like kinases, specifically AtBIN2^28^, whereas AtBASL recruits several members of the mitogen-activated protein kinase (MAPK) signalling cascade to its polarity crescent^30^. Polarisation of AtBASL, AtBRXL2 and AtPOLAR proteins is highly interdependent in *A. thaliana*, and the POLAR family seems largely redundant^26,28^.

However, in grasses, the *POLAR* gene family seems expanded, suggesting a functional divergence linked to the development of grass-specific epidermal features such as paracytic SCs^31^. Phylogenetic gene analysis revealed three major clades of at least five *POLAR* genes in grasses, with only one of them forming a clade with the *AtPOLAR* family, suggesting functional diversification not only between *Brassicaceae* and *Poaceae*, but also within the grass POLAR family (Figure 1B). Furthermore, the important polariser of the *A. thaliana* stomatal lineage BASL seems to be specific to eudicots and largely absent in grasses^31,32^, suggesting that grasses may employ different mechanisms for establishing polarity in epidermal lineages.

Indeed, BdPOLAR was identified as a regulator of SMC division in *B. distachyon*^33^. Single mutations in *BdPOLAR* resulted in defective SC divisions, with oblique and misoriented division planes, leading to missing or malformed SCs (Figure 1C)^33^. Reporter lines revealed that BdPOLAR forms a distal polarity domain (Figure 1D), which is opposed to the proximal polarity domain at the GMC/SMC interface of BdPANGLOSS1 (BdPAN1), a known SMC polarity factor in grasses^34,35^. These reciprocally opposing domains were shown to specify cortical division sites and control nuclear migration, coordinating the longitudinal ACD of the SMC, which is crucial for grass stomatal morphology and function^33,36^. However, *bdpolar-1* mutants displayed a moderate phenotype with only 25 - 40% of SCs affected^33^. This suggests potential residual redundancy in the grass POLAR gene family and, therefore, prompted us to analyse the role of *BdPOLAR’s* closest homolog that showed epidermis-specific expression, *BdPOLAR*-like1 (*BdPL1*) gene (Figure S1A).

Here, we identified BdPL1 as an intrinsically polarised polarity factor that operates across all epidermal cell files during the stage 2 patterning divisions in the developing *B. distachyon* epidermis. Fusion-protein reporter lines showed that BdPL1 polarised along the apical-basal axis before transverse ACDs, marking the future basal daughter cell and potentially positioning the cortical division sites. Much like for BdPOLAR, BdPL1’s polarisation was transient, and BdPL1 rapidly dissociated from its basal polarisation domain at the onset of division. Loss of *BdPL1* disrupted post-division cell size asymmetry, causing pavement cell clusters, impaired between-file coordination of epidermal patterning, and transversely split SCs. Ectopic BdPL1 expression caused excessive transverse divisions, and abolishing polarization of BdPL1 disrupted its function. Ectopic expression of BdPOLAR and BdPL1 under their reciprocal promoters showed that BdPL1 is intrinsically polarised independently of cellular context, whereas BdPOLAR is polarised in a context-dependent manner. Finally, we demonstrated that the known regulators of leaf epidermal patterning in grasses BdYODA1 (BdYDA1) and BdBRX-solo^37,38^, both polarise at the basal membrane post-division and independently of BdPL1. We propose a model in which pre-division and post-division polarity modules act sequentially and independently to robustly pattern the grass leaf epidermis and form the synapomorphic long-short epidermal pattern of grasses.

## Results

### *BdPL1* is required for the transverse asymmetric patterning divisions

To determine the biological function of *BdPL1*, we generated different loss-of-function alleles using CRISPR/ Cas9 (Figure 1E). The *bdpl1* mutants showed aberrant transverse asymmetric patterning divisions in all cell files (i.e. stomatal and hair cell files; Figures 1F, 1G, S1B-C). Instead of generating the stereotypical pattern with smaller apical cells and larger basal cells at stage 2 (Figure 1F), *bdpl1* mutants displayed misoriented, sometimes oblique division planes with weak or even inverted size asymmetries post-divisions (Figures 1G, S1B-C). In wild type (WT), robust cell size asymmetry within files ensured that small apical daughter cells are neighbouring bigger basal daughter cells from the adjacent cell files to avoid three-way junctions next to GMCs or hair cell precursors. In *bdpl1*, the misoriented division planes distorted medio-lateral patterning, mispositioning GMCs next to three-way junctions, which disrupted SC formation and produced bisected SCs, with “half” of the SC recruited from the upper SMC and the other “half” from the lower SMC (Figures 1G, S1B-C). This SC-defective phenotype, however, was distinct from the *bdpolar*-induced SC phenotype, where SMCs failed to polarise and divided obliquely (Figure 1C)^33^.

Together, this suggests that BdPL1 functions during the early transverse patterning division in all cell files, which is distinct from BdPOLAR’s SC-lineage-specific function.

### BdPL1 is polarised pre-division

To examine where and when BdPL1 is expressed, we generated *B. distachyon* lines expressing a transcriptional (*BdPL1p:mCit-mCit*^*NLS*^) or a translational (*BdPL1p:BdPL1-mVenus*) reporter gene. Transcriptional reporters showed that the *BdPL1* promoter activity started in late stage 1 protodermal cells, and peaked just before the transverse ACDs in all cell files in stage 2 (Figure 2A). Promoter activity then decreased until stage 4, where *de novo* expression was again detected in GMCs (Figure 2A). Fluorescent protein signal persisted in the young and maturing GCs during stages 5 and 6, but whether this signal was due to maintained expression or fluorescent protein stability was unclear.

**Figure 2.**
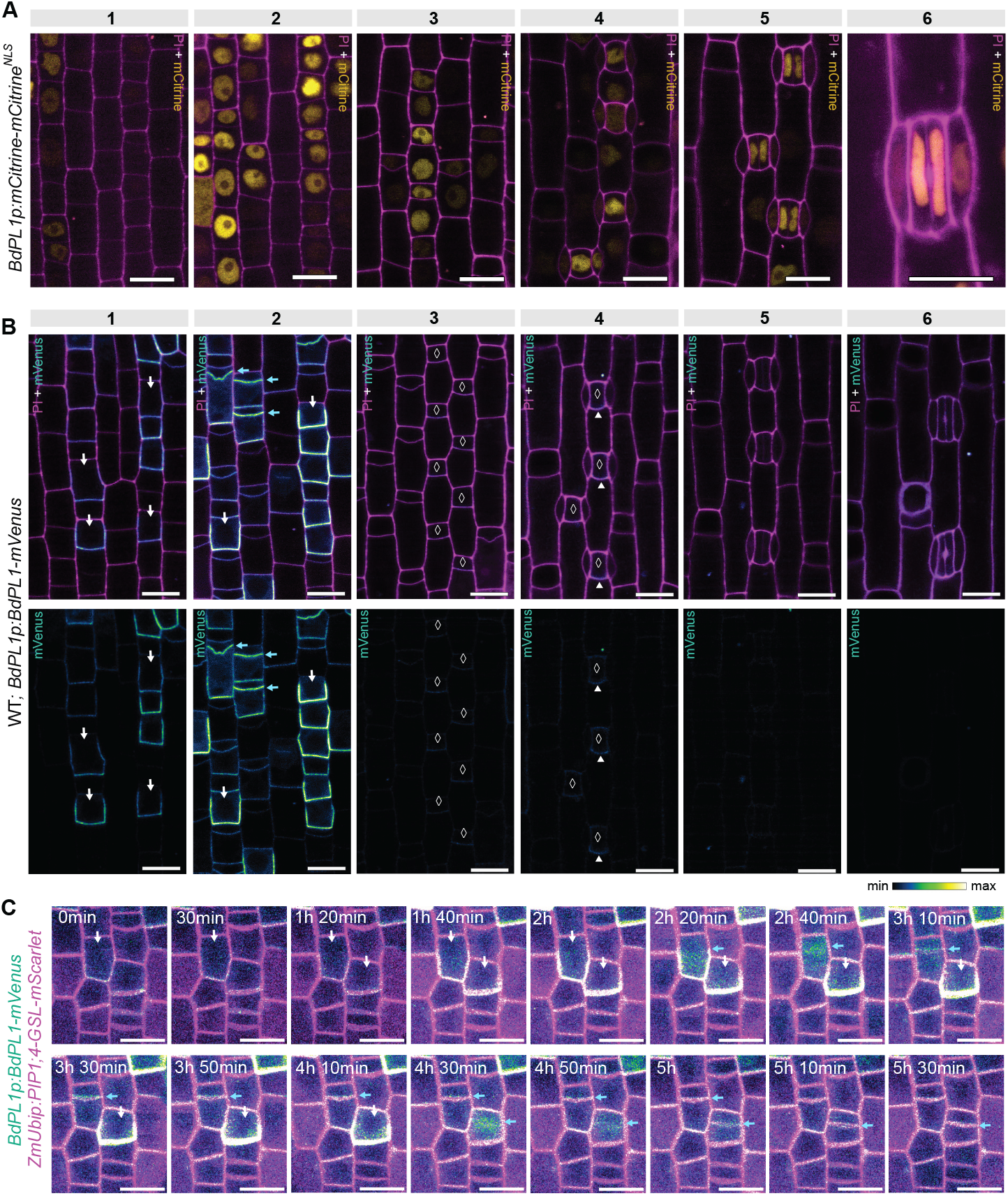
Polar localisation of BdPL1 precedes transverse patterning divisions. **(A)** Expression pattern of the transcriptional *BdPL1p:mCitrine-mCitrine*^*NLS*^ reporter in the developing leaf epidermis. Nuclear mCitrine signal is shown in yellow and propidium iodide (PI)-stained cell walls in magenta. Developmental stages are indicated. **(B)** Expression pattern of the translational *BdPL1p:BdPL1-mVenus* reporter in the developing leaf epidermis. Upper panels show merged signals of polarised BdPL1-mVenus (cyan) and PI-stained cell walls (magenta), lower panels show only the BdPL1-mVenus signal (“Green Fire Blue” heatmap). White arrows indicate pre-division polarisation of BdPL1-mVenus, cyan arrows point at post-division relocalisation of BdPL1-mVenus, white diamonds indicate GMCs, white arrowheads point at weak polarisation of BdPL1-mVenus in GMCs. Developmental stages are indicated. **(C)** Time-lapse imaging of the translational reporter *BdPL1p:BdPL1-mVenus* (cyan) introgressed into the plasma-membrane marker line *ZmUBIp:PIP1;4-mScarlet* (magenta) during transverse patterning divisions. White arrows indicate polarisation of BdPL1-mVenus, cyan arrows indicate relocalisation of BdPL1-mVenus. Time points are indicated. Scale bars, 10 µm.

In contrast, the translational BdPL1-mVenus fusion protein showed strong and almost exclusive expression during stage 2 transverse ACDs. It appeared to be polarised at the basal and most of the lateral cell walls just before stage 2 ACDs (Figure 2B). Strikingly, BdPL1 localisation seemed to stop just below the future division plane, demarcating the future basal daughter cell (i.e. future pavement cell). This “distal” polarisation (i.e., distal to the smaller daughter cell) of BdPL1 is highly reminiscent of the distal polarisation pattern of BdPOLAR, yet following the apical-basal instead of the medio-lateral polarization field^33^. After stage 2 ACDs, BdPL1-mVenus was hardly detected, apart from a very weak, basally polarised signal in stage 4 GMCs (Figure 2B). Although the difference between the transcriptional and translational reporters could reflect greater stability of the free fluorescent protein, this also may suggest that even though the *BdPL1* promoter was active in the GC lineage (Figure 2A), the protein was not translated or was rapidly turned over (Figures 2B, S2A). Finally, we found that YFP-tagged BdPL1 was again expressed and polarly localised after the stage 2 ACDs in epidermal cell files above leaf veins, where late transverse ACDs generated silica cell precursors (Figures S2B-D). This strongly suggested that BdPL1 was stabilised and basally polarised in all cells undergoing a transverse ACD in the developing grass leaf epidermis.

BdPL1’s basal localisation seemed to be of a transient nature only, as the polarised BdPL1-mVenus signal quickly disappeared postmitotically and seemed to re-localise to the newly formed plasma membrane (PM) between apical and basal daughter cells (Figure 2B). To observe the dynamics of this re-localisation, we performed time-lapse imaging of dividing stage 2 epidermal cells with the translational reporter line introgressed into a plasma membrane marker line (*ZmUbi1p:AtPIP1;4-mScarlett*). Indeed, upon strongly polarised basal localisation, BdPL1-mVenus started dissociating from the basal PM at the onset of mitosis and coalesced at the newly forming transverse PM (Figure 2C). This process was rapid, suggesting re-localisation of existing protein rather than *de novo* translation.

In conclusion, BdPL1 transiently polarises before stage 2 transverse ACD along the apical-basal axis, marking the large, basal daughter cell in all cell files.

### BdPL1 polarisation is required for protein function

To confirm that the function of the BdPL1 protein is not disturbed by a fused fluorescent tag, we transformed *bdpl1-1* mutants with *BdPL1p:BdPL1-mCitrine* or *BdPL1p:mCitrine-BdPL1* reporter constructs. Both BdPL1-mCitrine and mCitrine-BdPL1 displayed the previously observed, basal polarisation pattern and seemed to be able to complement the *bdpl1* phenotype and restore the physical asymmetry of the transverse ACDs (Figures 3A, S2A and S3A).

**Figure 3.**
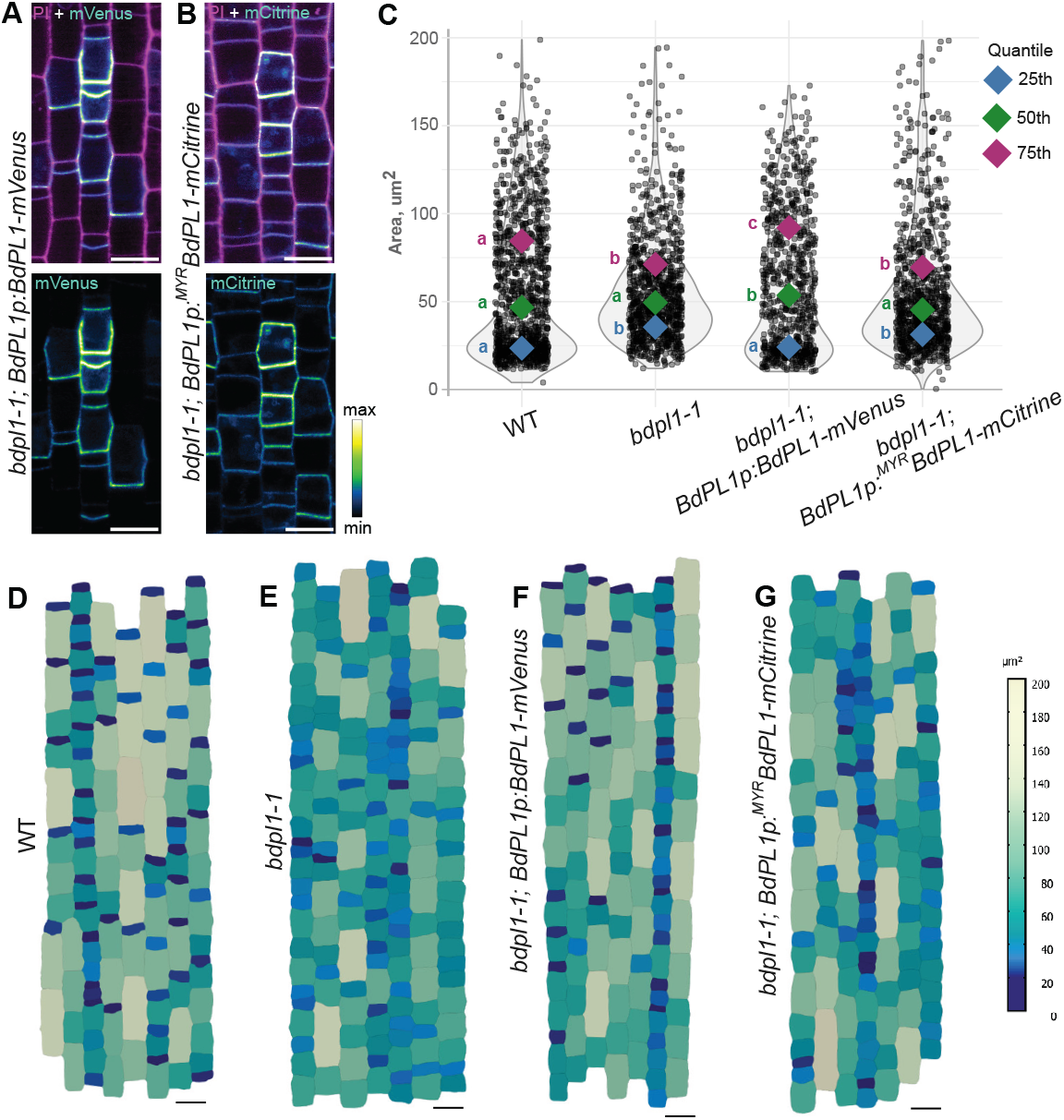
*BdPL1* enforces physical cell size asymmetry after the division. **(A-B)** Developmental zone of *bdpl1-1* mutant epidermis expressing *BdPL1p:BdPL1-mVenus* (A) or *BdPL1p:*^*MYR*^*BdPL1-mCitrine* (B). Upper panels show merged mVenus/mCitrine (cyan) signals and PI-stained cell walls (magenta), lower panels show only the mVenus/ mCitrine signal (“Green Fire Blue” heatmap). Scale bars, 10 µm. **(C)** Quantile regression analysis of cell areas post-transverse ACDs in the developmental zone of wild-type (WT), *bdpl1-1*, and *bdpl1-1* expressing a functional (*BdPL1p:BdPL1-mCitrine;bdpl1-1*) and non-functional (*BdPL1p:*^*MYR*^*BdPL1-mCitrine;bdpl1-1*) reporter construct. Violin plots show distributions of individual cell areas (grey dots); coloured diamonds indicate 25th (blue), 50th (median, green), and 75th (purple) quantiles. Letters indicate statistical groupings based on quantile regression with WT as the intercept (p-values listed in Table S1). **(D-G)** Segmented images of developing leaf epidermis post-transverse ACDs for WT (D), *bdpl1-1* (E), *BdPL1p:BdPL1-mCitrine* in *bdpl1-1* (F), and *BdPL1p:*^*MYR*^*BdPL1-mCitrine* in *bdpl1-1* (G). Cells are colour-coded based on the area (μm^2^) according to the scale (right). Scale bars, 10 µm.

To determine whether BdPL1’s polarisation is required for its role in ACDs, we generated lines disrupting BdPL1 polarisation by adding a short myristoylation acceptor peptide to the BdPL1-mCitrine fusion protein (*BdPL1p:*^*MYR*^*BdPL1-mCitrine*), which constitutively tethers the protein to the PM and disrupts polarised localisation^26,33^. Indeed, the ^MYR^BdPL1-mCitrine fusion protein was uniformly distributed across the PM (Figures 3B, S3B) and seemed unable to rescue the *bdpl1* phenotype (Figures 3B, S3B).

To quantitatively characterise complemented and non-complemented *bdpl1* mutant lines, we imaged stage 2 leaf epidermal zones in WT, *bdpl1-1*, and *bdpl1-1* expressing BdPL1-mCitrine or ^MYR^BdPL1-mCitrine. We segmented these images using MorphoGraphX^39^ and quantified cell area (Figures 3D-G). After transverse ACDs, WT and *bdpl1-1* expressing BdPL1-mCitrine showed clear physical asymmetry in both stomatal and non-stomatal cell files with small specialised cell precursors (GMCs or prickle hair cell precursors) apically and large pavement cell precursors basally (Figures 3D, 3F). In contrast, *bdpl1-1* mutants and *bdpl1-1* expressing ^MYR^BdPL1-mCitrine showed defective cell-size asymmetries (Figures 3E, 3G).

Quantification of cell area distribution in these four genotypes showed a narrower distribution in genotypes without functional or polarised BdPL1 (Figure 3C). We built quantile regression models, where we estimated the effects of genotype on cell area at the 25^th^, 50^th^ (median), and 75^th^ percentiles (Figures 3C, S3C). WT was used as the reference genotype, so the genotype coefficients for the mutant and rescue lines represent differences relative to WT. At the low end of the distribution (25^th^ percentile), complemented lines do not differ from the WT, while *bdpl1-1* and myristoylated lines showed higher cell area values. At the median, no statistically significant differences were found. However, at the upper end (75^th^ percentile), both *bdpl1-1* and myristoylated lines showed significantly lower cell area values (Table S1). Together, compressed distributions of *bdpl1-1* and myristoylated lines indicated disrupted post-division cell size asymmetry, while WT and complemented lines displayed distinct populations of smaller cells and wider range of cell areas between the median and 75^th^ percentile. Yet, even though the cell size distribution was affected, the cell density at the stage 2 did not show any significant changes (Figure S3D), indicating that cell division capacity was not impaired.

Overall, these results strongly suggested that basal polarisation is essential for BdPL1’s function and ability to enforce the physical asymmetry during cell division.

### BdPL1 is intrinsically polarised

Polarised BdPL1 is required for properly oriented patterning divisions. Yet, if BdPL1 is intrinsically polarised or if it requires additional stage-specific polarity cues remains unknown. Much like BdPOLAR, which requires BdPAN1 to be polarised along the medio-lateral axis in SMCs^33^, basal BdPL1 polarisation might depend on other polarised agonists and/or antagonists that are co-expressed in stage 1 cells.

To test this, we swapped the expression domains of BdPOLAR and BdPL1, which are medio-laterally polarized in SMCs and apical-basally polarized in stage 1 protodermal cells, respectively (Figure 4A). We generated promoter-swap lines that express BdPL1 in stage 3 SMCs (*BdPOLARp:BdPL1-mVenus*) and BdPOLAR in stage 1 protodermal cells (i.e. *BdPL1p:BdPOLAR-mCitrine*). We then imaged these reporter lines and compared them to their endogenous promoter-driven polarity profiles (Figures 4D-G).

**Figure 4.**
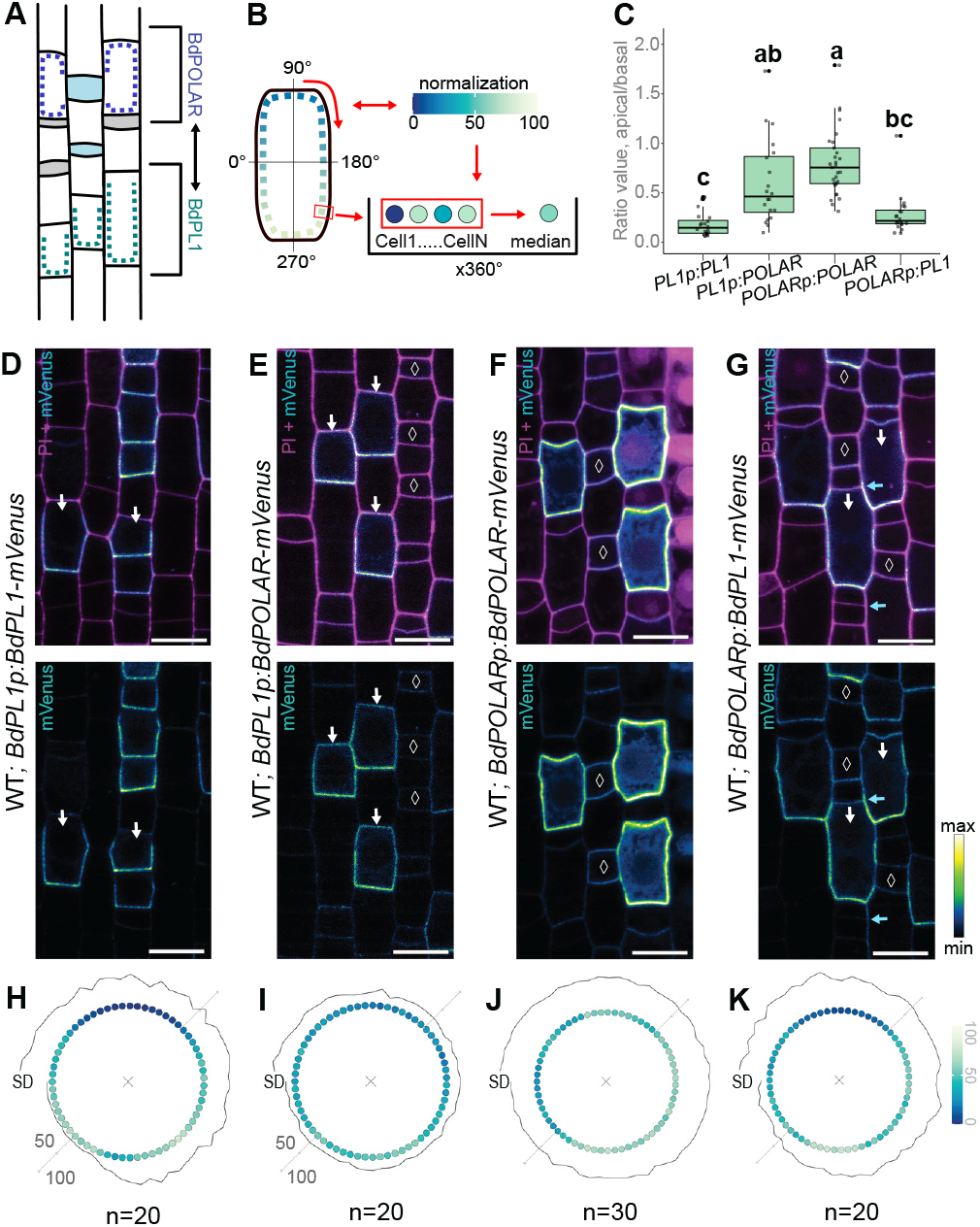
BdPL1 is intrinsically polarised independently of cellular context. **(A)** Schematic illustration of BdPL1 (teal) and BdPOLAR (purple) expression domains and localisation in developing epidermis. **(B)** Diagram of the fluorescence intensity quantification method used to calculate polarity profiles. Fluorescence along the cell cortex was measured from 0° to 360° in a clockwise direction and normalised for each cell (0-100%). Median values of normalised intensity values across all analysed cells were used to generate polarity profiles. **(C)** Quantification of apical-basal fluorescence ratios for the indicated reporter lines. Each box represents pooled values from 20-30 cells per genotype. Boxplots show the median and interquartile range, whiskers represent standard deviation (SD). Letters indicate statistically distinct groups based on one-way ANOVA followed by Tukey’s post-hoc test (p-value < 0.05). **(D-G)** Representative confocal images of the developing epidermis expressing *BdPL1p:B-dPL1-mVenus* (D), *BdPL1p:BdPOLAR-mVenus* (E), *BdPOLARp:BdPOLAR-mVenus* (F), *BdPOLARp:BdPL1-mVenus* (G) in WT. Upper panels show merged mVenus/mCitrine (cyan) signals and propidium iodide (PI)-stained cell walls (magenta), lower panels show only mVenus/ mCitrine signal (“Green Fire Blue” heatmap). White arrows point to cells polarised along the apical-basal axis, cyan arrows point to additional transverse divisions, white diamonds indicate GMCs. Scale bars, 10 µm.

We used POME^40^ to track if and how these polarity proteins polarise when placed in a distinct cellular context. To ensure that fluorescence intensity was not affected by neighbouring cells, we manually selected spatially isolated cells for each line (Figures S4A-D). We adjusted POME so that the fluorescence intensity measurement along the cell perimeter always started on the left side of the cell along the horizontal middle axis, which corresponded to 0º (Figure 4B, https://github.com/raissig-lindner-lab/Korostel-eva-et-al_2026_BdPL1). All intensity measurements were taken in a clockwise direction, allowing for the comparison of the absolute and relative intensities along the cell membrane for different cell types. Since BdPOLAR in SMCs is polarised along the medio-lateral axis based on the position of the GMC, we mirrored SMCs located on the left side of the GMC, so that all the images of polarised BdPOLAR or ectopically expressed BdPL1 in SMCs are oriented in the same direction (i.e., right-sided; Figure S4C). Based on these measurements, we generated polarity domain profiles for each construct (Figures 4H-4K), where absolute values were normalised to a 100%-scale for each cell and then the median across all cells in the sample was calculated for each measured angle, creating a median, relative polarity domain profile that we visualised in “solar flare” plots (Figures 4H-4K).

BdPOLAR misexpressed in stage 1 protodermal cells was able to weakly polarise before transverse asymmetric divisions along the apical-basal axis, yet much less pronounced than BdPL1 under its own promoter (Figures 4E and 4D). Indeed, POME quantification of relative fluorescence intensity along the cell outlines showed that BdPOLAR lacked the strong asymmetry in distribution compared to BdPL1 (Figures 4I and 4H). Even though BdPOLAR still showed some preference towards the basal part of the cell, it also showed clear localisation at the apical part of the cell (Figures 4E and 4I). In contrast, the BdPL1 protein had almost no signal at the apical part of the cell membrane (Figures 4D and 4H). To further validate this observation, we calculated the signal intensity ratio between apical (i.e. 90º) and basal (i.e. 270º) domains. While BdPL1 was strongly polarised in its own context (median of less than ∼0.2, all ratio values below 0.5, indicating strong polarisation), polarisation is rather weak for BdPOLAR in stage 1 protodermal cells, with a median below 0.5, but very high dispersion of values (Figure 4C).

In contrast, BdPL1 expressed under the *BdPOLAR* promoter maintained its “intrinsic” basal polarisation, and did not follow medio-lateral polarity field of SMCs (Figures 4G, 4K). BdPL1 expressed in SMCs also showed very low apical-basal ratio values comparable to those observed when BdPL1 was expressed in its native context (Figure 4C).

In conclusion, BdPL1 maintained a stable apical-basal polarisation irrespective of cellular and developmental contexts. This indicated that, unlike BdPOLAR, BdPL1 was intrinsically polarised and followed the longitudinal polarisation axis irrespective of cellular context.

### BdPL1 enforces apical-basal polarity and promotes ectopic transverse divisions

BdPL1 and BdPOLAR are unstructured, seemingly disordered proteins with 53% similarity, which is consistent with characteristics of scaffolding proteins^41,42^ (Figure S5A). To functionally test if this degree of homology between BdPL1 and BdPOLAR is sufficient to partially complement each other’s absence, we transformed the promoter swap lines in their respective mutant backgrounds *(BdPL1:BdPOLAR-mVenus* in *bdpl1-1* and *BdPOLARp:BdPL1-mVenus* in *bdpolar-1*; Figures 5A and 5B).

**Figure 5.**
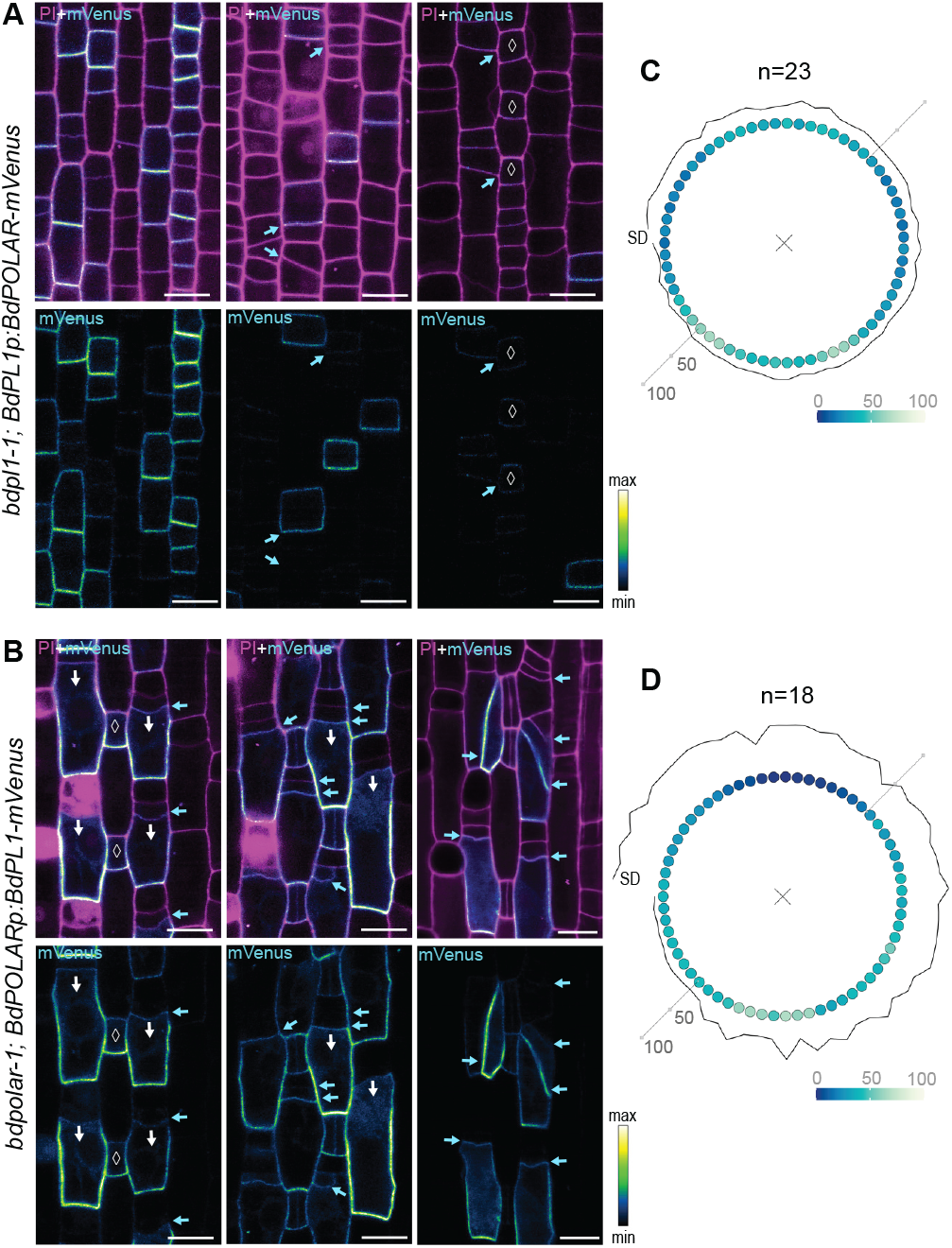
*BdPL1* and *BdPOLAR* cannot functionally complement each other. **(A-B)** Confocal images of *BdPL1p:BdPOLAR-mVenus* in the *bdpl1-1* background (A) and *BdPOLARp:BdPL1-mVenus* in the *bdpolar-1* background (B). Upper panels show merged mVenus (cyan) signal and propidium iodide (PI)-stained cell walls (magenta), lower panels show only the mVenus signal (“Green Fire Blue” heatmap). White arrows indicate polarisation of BdPL1-mVenus, cyan arrows point at additional and misoriented divisions, white diamonds indicate guard mother cells (GMCs). Scale bars, 10 µm. **(C-D)** “Solar flare” polarisation profiles of *BdPL1p:BdPOLAR-mVenus* in *bdpl1-1* (C) and *BdPOLARp:BdPL1-mVenus* in *bdpolar-1* (D). Radial plots show median normalised fluorescence intensity (colour-coded range of 0-100%, bottom right) is plotted as a function of angular position (0°-360°). Standard deviation (SD) of raw values is indicated as a line outside of the polarity profile, n - number of individual cells.

While the *BdPL1:BdPOLAR-mVenus* construct showed weak apical-basal polarisation in WT (Figures 4E and 4I), it appeared completely unpolarised in the *bdpl1-1* background, with almost equal fluorescence signal intensities at both the apical and basal cell membrane (Figures 5A and 5C, S5B and S5F). Furthermore, unpolarized BdPOLAR in stage 1 protodermal cells seemed unable to (even partially) rescue the *bdpl1* developmental phenotype (Figure 5A).

Strikingly, BdPL1 polarisation in *bdpolar-1* SMCs was even more pronounced along the apical-basal axis than in WT SMCs and caused excessive additional transverse divisions in SMCs (Figures 5B, 5D, S5C and S5F). Similarly, BdPL1 polarisation and excessive transverse divisions were also pronounced in the *bdpan1* mutant background (Figure S5E). Additional transverse divisions were also promoted in the wild-type background, but to a lesser extent (Figure S5D). As a result, misexpressing BdPL1 in SMCs enforced apical-basal polarity, disrupted the longitudinal division plane of SMCs, resulted in oblique and ectopic divisions disrupting SC formation, and, therefore, failed to rescue the *bdpolar-1* mutant phenotype (Figures 5B, S5D and S5E).

Overall, this suggested BdPL1 polarity to be instructive, as it could override the medio-lateral polarisation of SMCs and induce ectopic transverse or oblique divisions.

### *bdpl1* mutants display cell patterning defects in mature leaves

BdPL1 is essential for the postmitotic size asymmetry of the stage 2 epidermal development. In mature *bdpl1* mutant leaves, however, early disruption of postmitotic size asymmetry leads to relatively mild mature patterning defects; *bdpl1* showed primarily within-file pavement cell clusters and between-file hair cell clusters, which are distinct from within-file hair cell and stomatal clusters found in the *bdyda1-1* mutant (Figures 6A-C)^37^. In *bdpl1*, however, hair cell precursors and GMCs were often placed next to the three-way junctions of neighbouring cells, leading to lateral hair cell clusters and bisected SCs, respectively (Figures 6B, S6A and S6B). These abnormalities indicate that defective cell size asymmetry in *bdpl1* affected medio-lateral patterning, where each specialised cell type (i.e. smaller apical cell) is positioned centrally between pavement cells rather than adjacent to pavement cell junctions (Figures 6A and 6B).

**Figure 6.**
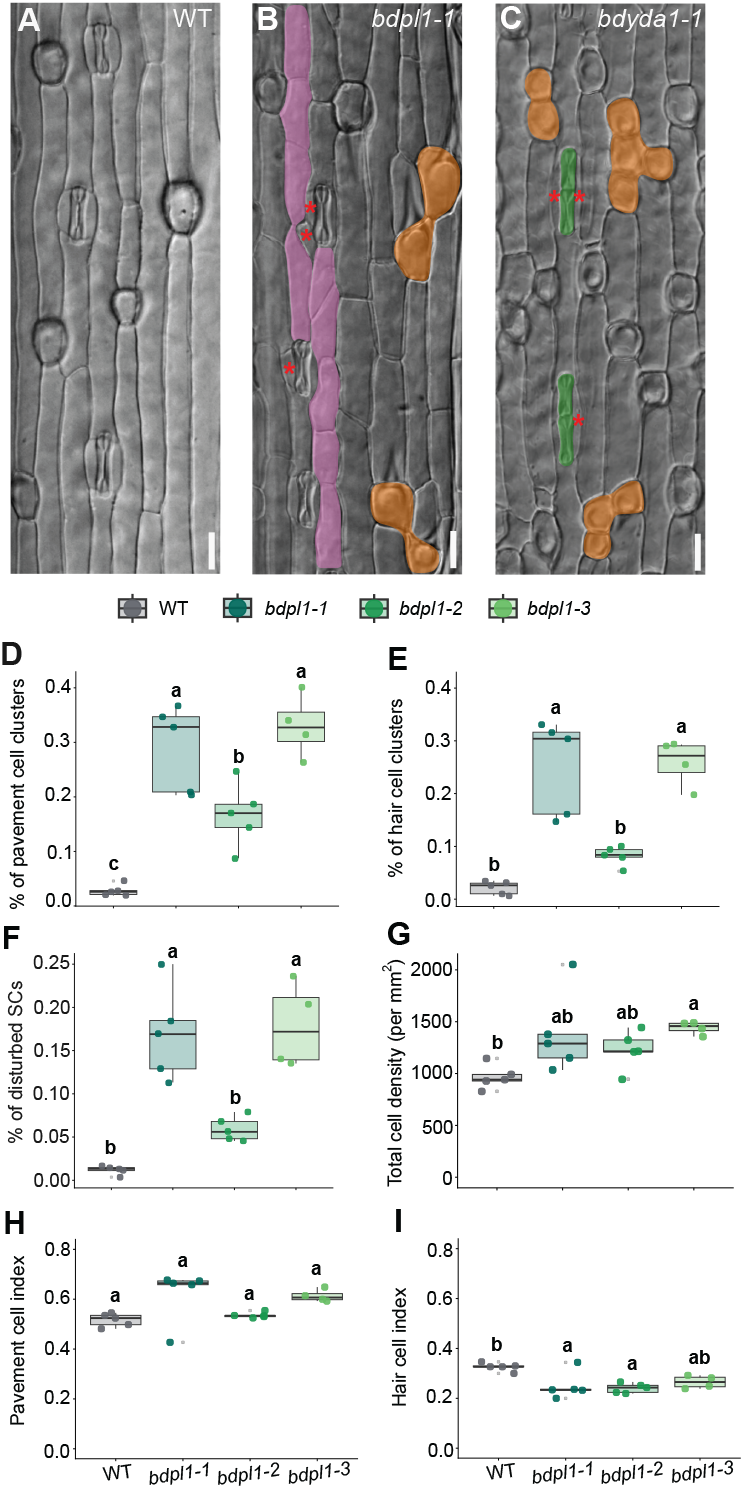
Loss of *BdPL1* disrupts the patterning of the mature leaf epidermis. **(A-C)** Differential interference contrast (DIC) images of the mature adaxial leaf epidermis of 21-day-old, soil-grown wild-type (WT) (A), *bdpl1-1* (B), and *bdyda1-1* (C) plants. Pavement cell clusters are false-coloured in pink, hair cell clusters are false-coloured in orange and disturbed SCs are indicated with red asterisks. Scale bars, 20 µm. **(D-F)** Quantification of proportion of clustered pavement cells (D), pseudo-clustered (i.e. between files) hair cells (E) and disturbed subsidiary cells (F) in wild type (WT) and three *bdpl1* alleles. (G) Total epidermal cell density per mm^2^ across three *bdpl1* alleles and WT. **(H-I)** Quantification of pavement cell index (H) and hair cell index (I) in WT and three *bdpl1* alleles. Measuring and quantification details can be found in STAR Methods, counting data can be found in Table S3. Boxes represent median and interquartile ranges; whiskers represent standard deviation (SD); dots indicate biological replicates; letters indicate statistically distinct groups determined by one-way ANOVA followed by Tukey’s post hoc test (p-value < 0.05).

Within-file stomatal clusters, however, were extremely rare; they appeared in several individuals, but did not form a consistent pattern across all three *bdpl1* mutant alleles (Figure S6A). Nonetheless, within-file pavement cell clusters and between-file hair cell clusters were consistently observed and affected 20-30% of all pavement and hair cells in *bdpl1* (Figures 6D and 6E), and approximately 17% of SCs appeared disturbed (Figure 6F). These numbers are lower for the *bdpl1-2* allele (Figures 6D-F), which has an in-frame deletion of 37 base pairs leading to a splice site removal, but seemingly not affecting correct translation of the second exon (Figure S6D). However, even in the hypomorphic *bdpl1-2* mutant, the patterning defects were obvious (Figure 6D-F).

Unlike in the developmental zone, the total cell density of the mature leaf epidermis was higher in *bdpl1* mutants (Figure 6G), which was reflected in a higher pavement cell index for *bdpl1-1* and *bdpl1-3* alleles (Figure 6H). However, it remains unclear whether increased cell density reflects a larger population of smaller pavement cells, impaired cell elongation, or both. Consistent with these findings, the hair cell index was reduced across all three *bdpl1* alleles (Figure 6I). Stomatal and silica cell indices were slightly lower in some alleles, though more variability was observed than for pavement and hair cell indices (Figures S6E-F).

To test residual genetic interaction of *BdPOLAR* and *BdPL1*, we generated the double knock-out mutant *bdpl1-4;bdpolar-4*. However, none of the three major phenotyping criteria (i.e. defective SCs, pavement or hair cell clusters) was significantly different between *bdpl1-1* and *bdpl1-4;bdpolar-4* mutants, but merely added *bdpolar*-like SMC division defects (Figures S6C, and S6G-S6I). These additive phenotypic defects indicate independent roles of *BdPL1* and *BdPOLAR* in epidermal development. Moreover, the relatively mild phenotype of *bdpl1* mutants in the mature grass leaf epidermis suggests that within-file asymmetric cell identity can be established independently of cell-size asymmetry indicating a post-mitotic compensation mechanism.

### Division of labour by a pre- and post-division polarity module

Considering the prominent developmental phenotype with disrupted stage 2 patterning divisions of *bdpl1* (Figure 1G), the rather mild patterning defects and particularly the absence of the “pearls-on-a-string” phenotype of specialized cell types (i.e. stomata, hair cells, silica cells) seen in *yda1* mutants was puzzling^37,38^. The triple MAP kinase *YDA* is known to enforce cell fate patterning asymmetry in the stomatal lineage in *A. thaliana* and in all epidermal lineages in grasses^37,38,43^. Yet, unlike BdPL1, BdYDA was suggested to polarise basally post-division rather than pre-division and showed wild-type-like ACDs at stage 2 protodermal cells^37,40^. Similarly, mutations in the putative polarisation factor HvBRX-solo (*cer-s*.*31*) in barley resulted in severe clustering of prickle hair and silica-cork cells^38^, which suggests that BRX-solo might act as a polarity scaffold in grass leaf epidermis development, much like the BRX family genes in the *A*.*thaliana* stomatal lineage^26,27,40^.

Consistent with previous reports^37,40^, we observed that *bdyda1-1* mutants performed transverse asymmetric patterning divisions normally, with clear post-division physical asymmetry. However, after stage 2, some cells underwent multiple rounds of additional transverse division, resulting in clustered small apical cells and, ultimately, to within-file clusters of stomatal precursors (Figure 7A).

**Figure 7.**
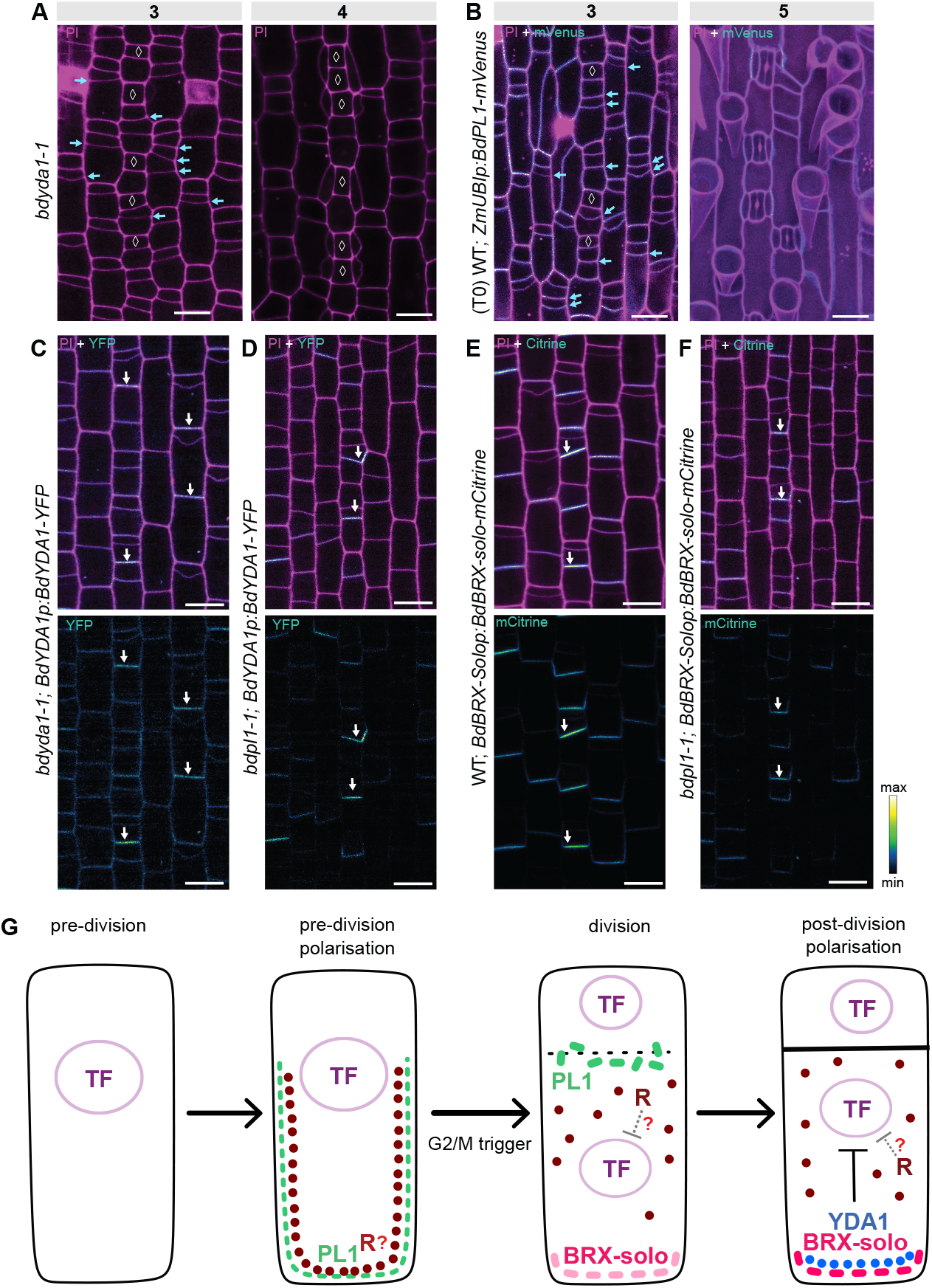
A premitotic and a postmitotic polarity module robustly pattern the grass leaf epidermis. **(A)** Confocal images of developing leaf epidermis in *bdyda1-1*. Cell walls stained with propidium iodide (PI, magenta). **(B)** Confocal images of developing leaf epidermis with ectopically expressed BdPL1-mVenus under the ubiquitous promoter (WT;*ZmUBIp:BdPL1-mVenus;* T0). Merged signals of mVenus (cyan) and PI-stained cell walls (magenta). Cyan arrows point at additional divisions, white diamonds indicate GMCs. Stages indicated. Scale bars, 10 µm. **(C-D)** *BdYDA1p:BdYDA1-YFP* expression in *bdyda1-1* (C) and *bdpl1-1* (D) backgrounds. **(E-F)** *BdBRX-solop:BdBRX-solo-mCitrine* expression in wild-type (E) and *bdpl1-1* (F) backgrounds. Upper panels show merged YFP/mCitrine signal (cyan) and PI-stained cell walls (magenta), lower panels show only YFP/mCitrine (“Green Fire Blue” heatmap). White arrows indicate post-division polarisation. Scale bars, 10 µm. **(G)** Proposed model illustrating the division of labour between pre- and post-division polarity modules. During pre-division polarisation, BdPL1 (green) marks the future basal daughter cell to establish physical asymmetry. Cell division initiation triggers the release of BdPL1 from the plasma membrane, and BdBRX-solo (pink) starts to accumulate at the basal part of the larger basal cell, i.e. pavement cell precursor. BdBRX-solo becomes enriched at the basal membrane and sequesters and polarized BdYDA1 together forming a post-division polarity module, which inhibits specialised cell identity in the larger basal daughter cell. The Transcription Factor (TF, purple) promotes specialised cell identity in the smaller apical daughter cell. We propose that a repressor of specialised cell fate (R) is sequestered by BdPL1, and not sequestering it in mutants leads to premature repression of specialised cell fate and the observed pavement cell clusters.

Similarly, overexpression of BdPL1 (*ZmUbi1p:BdPL1-mVenus*) caused additional transverse divisions in all cell files, forming stacks of small cells (Figure 7B, stage 3). Remarkably, these additional divisions did not affect future patterning to the expected extent: instead of “pearls-on-string” clusters of stomata or hair cells, specialised cells were separated by small pavement cells, maintaining correct patterning (Figure 7B, stage 5). This suggested that BdYDA1, potentially together with BdBRX-solo, can enforce cell fate asymmetry despite the ectopic divisions caused by overexpression of BdPL1.

To determine BdBRX-solo expression and polarisation in protodermal cells, we generated a reporter line for BdBRX-solo (*BdBRX-solop:BdBRX-solo-mCitrine*) and imaged developing leaf zones during transverse patterning divisions in the newly generated BdBRX-solo reporter and the previously published *BdYDA1* reporter line (*bdyda1-1*; *BdYDA1p:BdYDA1-YFP:Yt*) ^37^ (Figures 7C-F). BdYDA1 is broadly expressed and specifically polarised to the basal domain of pavement cell precursors, strictly after the transverse ACD (Figure 7C). Similarly, BdBRX-solo was polarised post-division during stage 2, showing weak polarisation right after the division before forming a strong and specific polarity domain later (Figure 7E). Notably, both proteins localised exclusively to the basal cell domain rather than the lateral walls, displaying a more restricted basal polarity than BdPL1.

To test whether the post-division polarity domains of BdYDA1 or BdBRX-solo depended on physical cell asymmetry or the pre-division polarity of BdPL1, we analysed their translational reporter lines in the *bdpl1* background. Both BdYDA1 and BdBRX-solo maintained their respective polarity domains and localised to the basal daughter cell post-division, even in the absence of physical asymmetry (Figures 7D, 7F and S7A). This suggested that the premitotic and postmitotic modules polarise independently of each other and control epidermal patterning and cell fate distribution sequentially.

Therefore, we propose a division-of-labour model (Figure 7G), where BdPL1 polarises pre-division, guiding the position of the division plane and establishing physical cell asymmetry across all epidermal files. This provides the foundation for epidermal patterning and prevents between-file cell clustering and three-way junctions next to hair cell precursors or GMCs that cause the transverse splitting of SCs. During the division, BdPL1 is released from the PM, which might allow BdBRX-solo to occupy the basal part of the dividing cell and recruit BdYDA1. BdYDA1 and BdBRX-solo fully polarise post-division, ensuring correct fate asymmetry and proper within-file patterning by repressing specialised cell fates in the basal daughter cell. These consecutive modules enforce the synapomorphic long-short pattern of grass leaf epidermis by ensuring robust medio-lateral and apical-basal cell patterning.

## Discussion

Our study identifies BdPL1 as an early-acting cellular polarity factor required for transverse asymmetric patterning divisions in the developing grass leaf epidermis. This function is distinct from that of its closest homolog, BdPOLAR, which polarises in SMCs and guides longitudinal asymmetric divisions^33^. While both BdPL1 and BdPOLAR form a polarity domain distal to the division plane, BdPL1 polarisation occurs broadly across all epidermal files and follows the apical-basal rather than the medio-lateral polarisation field. The physical cell-size asymmetry of the small apical and the large basal cell post-division seems to be necessary to enforce accurate between-file patterning and normal SC formation.

### How is transient BdPL1 polarisation established?

Polarised localisation can be established through different mechanisms. These include extrinsic, environmental cues such as light or gravity, organ-wide or tissue-wide axial polarity, or intrinsic cues such as membrane-associated polarity complexes that recruit other polarity factors as clients^3,44,45^. Together, all of these signals provide polarity axes that are subsequently interpreted and reinforced by transient polarity modules at the cellular level.

When ectopically expressed, BdPL1 was able to maintain its longitudinal polarisation independently of cellular context and developmental stage, suggesting that BdPL1 can either respond to extrinsic cues such as light or gravity, follow axial apical-basal tissue polarity of the growing leaf epidermis, rely on feedback from recruited effectors, or be the client of another, yet unknown, polarised protein. Alternatively, BdPL1 might act as a primary factor for establishing the longitudinal polarisation axis across the whole leaf epidermal tissue.

Nonetheless, the persistent polarisation of BdPL1 contrasts with that of BdPOLAR. In SMCs, BdPOLAR requires BdPAN1 to establish a medio-lateral, distal polarity domain and expands into the proximal domain in *bdpan1* mutants^33^. The ability of BdPL1 to maintain its apical-basal polarisation in SMCs, indicates that no stage-specific, reciprocally polarised protein is required. Apart from Bd-PAN1’s role in polarising BdPOLAR, there is little evidence in plants that two opposite polarity domains enforce each other’s reciprocal polarisation through reciprocal inhibition^33^. However, opposing polarity domains can be established independently, such as in *A*.*thaliana* root endodermis, where INFLORESCENCE AND ROOT APICES RECEPTOR KINASE (AtIRK) and KINASE ON THE INSIDE (AtKOIN) polarisation is generated through distinct sorting mechanisms^46^. A similar principle may apply in the stomatal lineage of *A. thaliana*, where AtOPL2 and the At-BASL-AtBRXf-AtPOLAR module occupy opposing cortical domains, but have not been shown to directly influence each other’s polar localisation^27^.

### How does BdPL1 implement physically asymmetric divisions?

BdPL1 expression and polarisation specifically occurred in all protodermal cells before transverse asymmetric patterning divisions in previously symmetrically dividing protodermal cells. Just before the asymmetric patterning division, BdPL1 was expressed and polarly localised, marking the future basal daughter cell and seemingly ending exactly where cortical division sites should be established (Figure 2B). This strongly suggests that BdPL1 is involved in breaking the symmetry of the previously symmetrically dividing protodermal cells. This symmetry breaking is required to establish the stereotypical pattern of the grass leaf epidermis, where small specialized cells are interspersed by long pavement cells within a file and flanked by pavement cells from adjacent files.

How BdPL1 implements ACDs remains unknown, but it might be associated with cortical microtubules (MTs) reorganisation within its polarity domain, which restricts preprophase band and cell plate formation to the apical region of the cell. In support of this idea, cortical MTs have been shown to accumulate in the apical parts of stomatal precursor cells in maize^47,48^. Additionally, BdPOLAR has been shown to affect the placement of cortical division sites, marked by BdTANGLED1 (BdTAN1), in SMCs outside of its polarity domain^33^. This organisation differs from *A*.*thaliana*, where the AtBASL-AtBRXf domain was depleted of stable MTs, while AtPOLAR co-localised with MTs outside of AtBASL-AtBRXf domain^49^. This suggests that the POLAR family function has diversified between Brassicaceae and grasses and indicates that the positioning of stable cortical MTs, and, consequently, preprophase band and cell plate formation outside of the POLAR family domain is a distinct feature in grasses.

Another distinctive feature of the grass POLAR proteins is their rapid dissociation and relocalisation to the new PM during mitosis. In *A. thaliana*, the stomatal polarity domains are inherited by the daughter cells and remain polarised post-division^25,26,29,49^. The molecular mechanism and functional relevance of BdPL1 relocalisation is unknown. It might simply follow general cellular vesicle trafficking during cytokinesis, which is primarily directed towards the forming cell plate^50^. However, BdPL1 relocalisation could deliver cellular effectors (i.e. polarity clients) to the newly formed cell plate to modulate division capacity or to reinforce cell fate asymmetry. Determining whether BdPL1 passively follows vesicle trafficking or actively re-distributes signalling components will require a comprehensive identification of BdPL1-associated effectors and their respective cellular dynamics.

### Is physical and fate asymmetry independent in grass leaf epidermal development?

BdPL1 polarised before the transverse patterning division, while BdYDA1 and BdBRX-solo became polarised only after the division. Even in *bdpl1*, both BdYDA1 and BdBRX-solo remained correctly polarised in basal daughter cells post-division, suggesting that the premitotic BdPL1 module and the postmitotic BdYDA-BdBRX-solo module function independently of one another.

Mutating either of the modules led to cell-type clusters, yet distinct cell types were affected. In *bdpl1*, pavement cells were clustered, whereas in *bdyda1-1, hvyda1* and *cer-s*.*31* (*hvbrx-solo*) specialised epidermal cells, like hair cells, stomatal complexes and silica cells, appeared in clusters^37,38^. This suggested that *yda1* and *brx-solo* mutants failed to repress specialised cell fate in the basal, large daughter cells, while *bdpl1* mutants repressed specialised epidermal cell identity prematurely.

We suggest that BdBRX-solo might act as a postmitotic scaffold sequestering BdYDA1 and enforcing asymmetric signalling to downregulate specialised epidermal cell identity in the basal daughter cell, similar to AtBASL’s role in sequestering the MAPK signalling module in *A. thaliana* SLGCs^30^. For BdPL1, we propose that beyond establishing physical cell size asymmetry, it also sequesters an effector that represses specialised cell fate. Polar localisation of BdPL1 enriches this repressor at the basal cell domain premitotically, ensuring asymmetric inheritance post-division (Figure 7G). Upon BdPL1’s release from the PM during mitosis, this putative repressor may act in concert with asymmetric MAPK signalling to robustly suppress specialised cell identity and enforce a non-specialised fate in the basal daughter cell post-division (Figure 7G).

The identity of this putative effector remains unknown. In *A. thaliana*, AtPOLAR scaffolds the GSK3 kinase AtBIN2 to the PM, where it phosphorylates and inhibits AtYDA, thus positively regulating AtSPCH activity before the meristemoid division^28,51,52^. At the onset of division, the protein phosphatase AtBSU1-like1 (AtBSL1) is recruited to the polarity crescent, releasing AtBIN2 and dephosphorylating AtYDA, which together robustly downregulate AtSPCH in the larger daughter cell (SLGC) post-division^53^. Thus, pre-division polarity establishment leads to post-division cell fate reinforcement through the dynamic relocalisation of signalling effectors^28,53^. In grasses, however, the putative polarity scaffolds BdPL1 and BdBRX-solo are polarised in a sequential, temporally separated manner, which omits the need for a regulator releasing effectors in a cell-cycle stage-specific manner, but instead might require a regulator that directly releases the BdPL1 polarity scaffold.

### Why do grasses employ a premitotic and a postmitotic polarity modules?

The divergence between eudicot and grass polarity systems likely reflects their contrasting epidermal tissue architectures, distinct patterning rules and divergent principal directions of growth in developing leaves. The development of the *A. thaliana* leaf epidermis allows flexible placement of stomata through iterative ACDs, and thus polarity must be repeatedly established considering primarily the cellular microenvironment of the surrounding cells^27,28,54^.

The grass leaf’s development along a strict acropetal gradient, with cell divisions restricted to a narrow, spatially defined zone near the shoot apex, might require more stringent programs that are specific to mitotic stages^7,8^. In the grass leaf epidermis, transverse asymmetric patterning divisions happen only once. Consequently, between-file cell positioning and cell fate distribution cannot be compensated through additional divisions. Therefore, BdPL1-mediated cell size-asymmetry might be required to maintain alternating, non-clustered patterning between adjacent cell files. Indeed, patterning divisions seem to occur in small, synchronised cell groups within a file, which might help to ensure the correct positioning of different cell types (i.e., apical specialised cell and basal pavement cell) between different cell files and prevent split SCs and between-file hair cell clusters. Finally, the postmitotic cell fate asymmetry module can compensate for sporadic division defects that can occur during patterning divisions and enforce “correct” patterning. Therefore, the temporal separation between the premitotic (BdPL1) and postmitotic (BdYDA1, BdBRX-solo) polarity modules may have evolved to ensure strict developmental patterning with minimal positional errors and thus robust cell-type distribution across the grass leaf epidermis.

## Supplemental Information

Document S1. Figures S1–S7, Tables S1-S2. Table S3.

## Acknowledgments

We would like to thank the research gardeners Sarah Dolder, Jasmin Sekulovski and Christopher Ball (University of Bern, Switzerland). We thank Guillaume Witz from the Data Science Lab (University of Bern, Switzerland) for assistance with adapting a published tool used in this study. We thank Jan Lohmann (COS Heidelberg, Germany) and Karin Schumacher (COS Heidelberg, Germany) for kindly providing plasmids, and Dominique Bergmann (Stanford University, United States) for seeds. Microscopy was performed on equipment supported by the Microscopy Imaging Centre (MIC) at the University of Bern. This work was supported by the Swiss National Science Foundation (SNSF) Grant 215019 (to M.T.R.) at the University of Bern.

## Author contributions

Conceptualisation: MTR; Data curation: ALK, KJ; Formal analysis: ALK, KJ; Funding acquisition: MTR; Investigation: ALK, KJ, DZ, IP, NG, EB; Methodology: ALK, PRD, BJ, HL, MTR; Project administration: MTR; Resources: MTR; Supervision: MTR; Validation: ALK, KJ, DZ; Visualisation: ALK, MTR; Writing – original draft: ALK, MTR; Writing – review & editing: all authors. All authors have read and approved the final manuscript.

## Declaration of interests

The authors declare no competing interests.

## Materials and Methods

### Key resources table

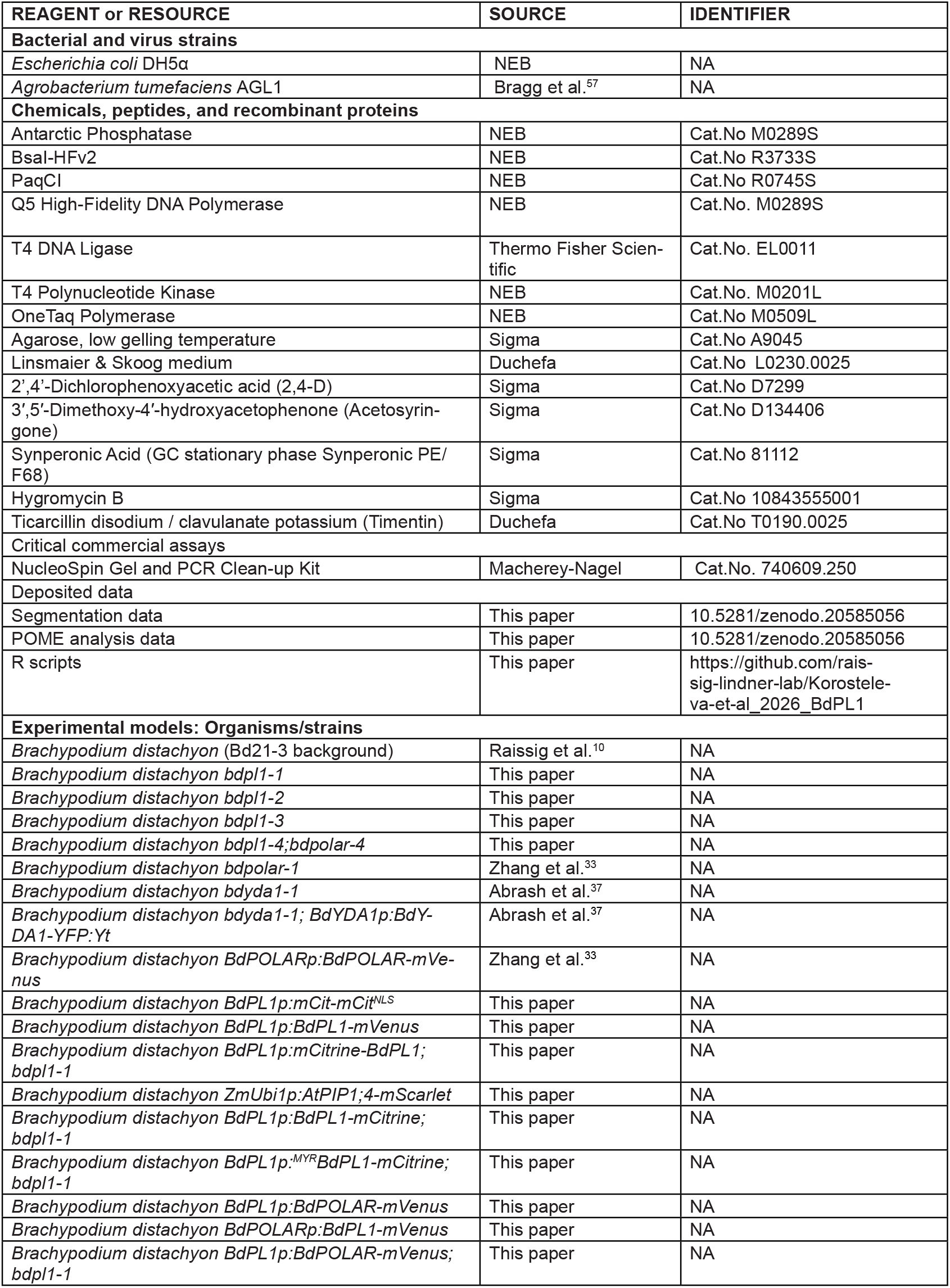
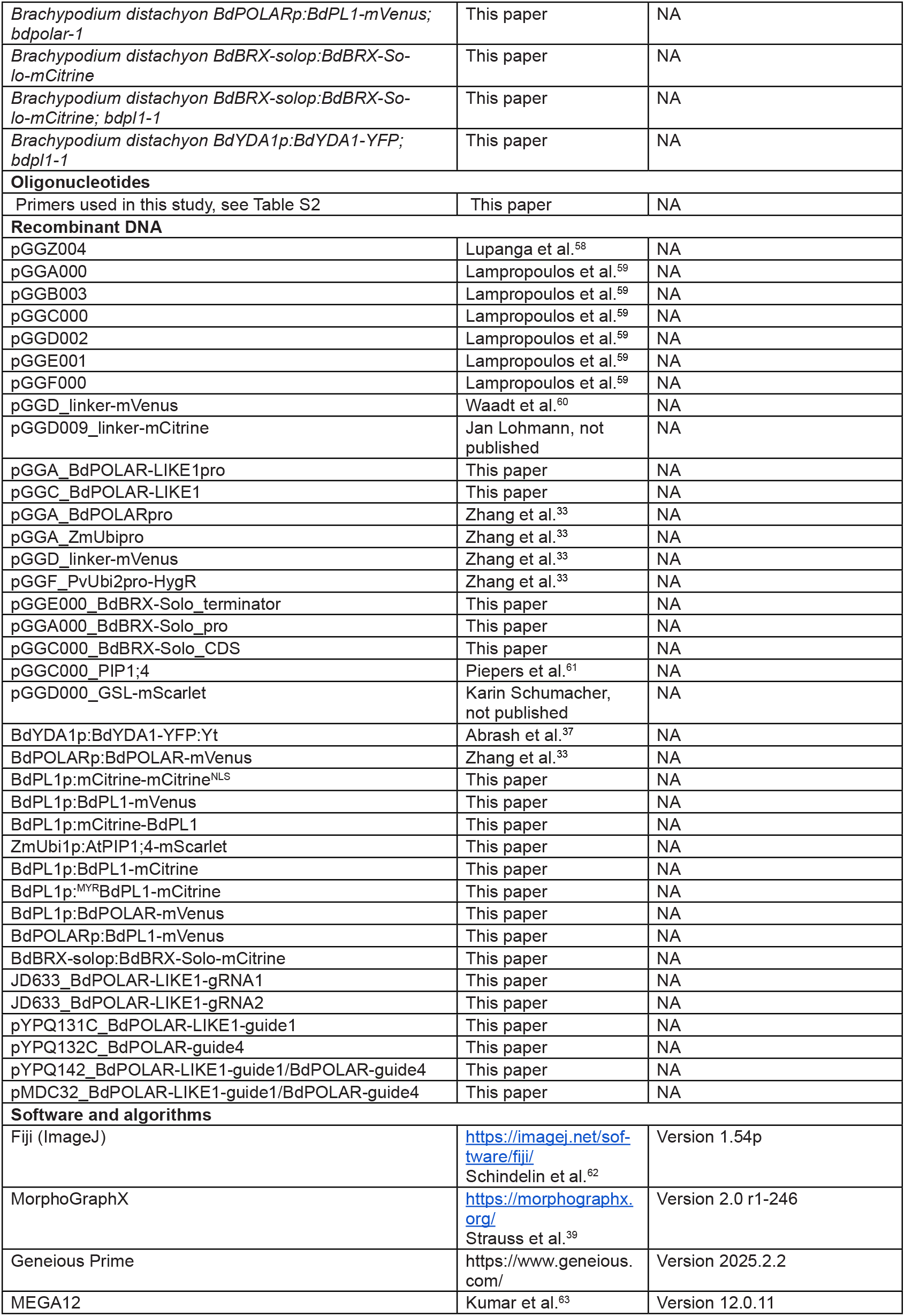
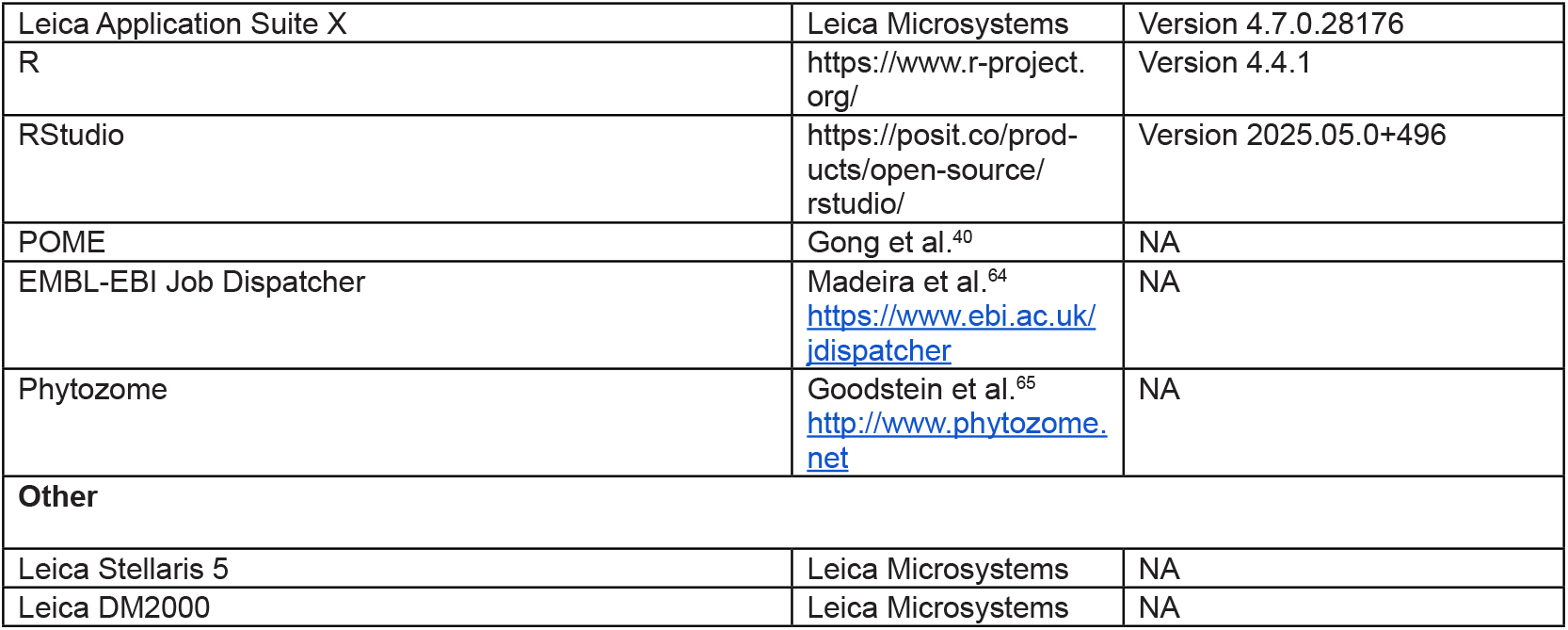

### Resource availability

#### Lead contact

All the materials generated in this manuscript are available upon request from the lead contact, Michael T. Raissig.

#### Materials availability

Plan materials and constructs generated in this study are available upon request from the lead contact.

#### Data and code availability

Microscopy images used for quantitative analysis are publicly available at Zenodo (10.5281/zenodo.20585056). All quantitative data generated and analysed in this study can be found in the supplementary materials. Adjusted POME FIJI macro and R scripts used to analyse the data in this study can be found on GitHub (https://github.com/rais-sig-lindner-lab/Korosteleva-et-al_2026_BdPL1). Any additional information in this paper is available from the lead contact upon request.

### Experimental model details

#### Plant material and growth conditions

*Brachypodium distachyon* Bd21-3 was used as wild type (WT) for all experiments. *bdpl1* mutants were obtained with CRISPR/Cas9 editing, and the *bdpl1-1* allele was used as the reference allele.

All seeds were sterilised for 5 min with 20% bleach (Roth), 0.1% Triton X-100 (Roth) and washed with sterile water. For genotyping, crossing and collection of the third mature leaf, the seeds were left in water and placed in the dark at 4°C for vernalization and stratification for at least two days. Afterwards, seeds were placed directly on soil in a growth chamber with a 18h light/6h dark cycle (PPFD 250-350 μmol m^-2^ s^-1^; day temperature 28°C, night temperature 22°C).

For confocal microscopy and time-lapse experiments, seeds were stratified on 1/2 Murashige and Skoog (1/2 MS) (Duchefa) plates with 1% agar (Duchefa), pH 5.7. After vernalization and stratification, these seeds were placed into a 28°C chamber with a 16h light/ 8h dark cycle and 110 μmol m^-2^ s^-1^ (as the unit of photosynthetically active photon flux density, PPFD) for 5-6 days until imaging of the second leaf’s developmental zone or for 3 days until the time-lapse experiment.

### Method details

#### Confocal microscopy

For imaging of the developmental zone, the second leaf from a 5-to 6-day-old plate-germinated seedling was gently pulled out and placed in a 15 μM Propidium Iodide solution (PI, 1:100 dilution from a 1.5 mM stock (Thermo Fisher Scientific)) for 5 minutes. Imaging was performed with a Leica Stellaris 5 confocal microscope with 10-20% laser intensity. YFP/mCitrine/mVenus and PI were simultaneously excited at 515 nm, and fluorescent emission was detected at 530-550 and 610-650 nm for respective channels. All confocal images, except time-lapse imaging, were made with 63x objective (Leica Microsystems, Product No. 11506398), 80% glycerol was used as immersion medium.

#### Sample preparation for live imaging

Time-lapse imaging was performed using the *BdPL1p:BdPL1-mVenus; ZmUBIp:PIP1;4-mScarlet* reporter line obtained by crossing the respective single transgenic lines. Three-day-old seedlings were dissected under a stereomicroscope on moist filter paper. The first leaf was pinched with a pin to prevent the seedling from moving, while the coleoptile and first leaf were carefully removed using fine tweezers, leaving the second developing leaf attached to the meristem and root for imaging.

Dissected seedlings were transferred to 60 mm Petri dishes containing 1/2MS medium with 2% agar (Duchefa). Roots were fixed with 3% low-melting-point agarose (Sigma/Merck) cooled to 35°C before application. Before imaging, a thin layer of water was added over the medium, and the water immersion 63x objective (Leica Microsystems, Product No. 11506362) was used. To avoid photobleaching, 5% laser intensity was used, with mVenus excited at 514nm and detected at 530-550nm, and mScarlet excited at 569nm and detected at 610-630nm.

#### Phylogenetic analysis and protein alignment

Protein sequences of *BdPOLAR* (BdiBd21-3.3G0715200) homologues were retrieved from Phytozome^65^ for *Arabidopsis thaliana* (TAIR10, genome ID: 167), *Brachypodium distachyon* (v1.2, genome ID: 537), *Hordeum vulgare* (r1, genome ID: 462), *Oryza sativa* (v7.0, genome ID: 323), *Zea mays* (RefGen_V4, genome ID: 493), *Solanum lycopersicum* (ITAG5.0, genome ID: 796) and *Populus trichocarpa* (v4.1, genome ID: 533). Amino acid sequences were aligned in MEGA12^63^ using MUSCLE. The resulting alignment was used to construct a phylogenetic tree in MEGA12 using the maximum likelihood method with Jones-Taylor-Thornton (JTT) amino-acid substitution model. Protein sequence similarity for *BdPOLAR* (BdiBd21-3.3G0715200) and *BdPL1* (BdiBd21-3.1G0621800) was calculated with EBLOSUM62 matrix score using EMBOSS Water at EMBL-EBI Job Dispatcher^64^ (https://www.ebi.ac.uk/jdispatcher).

#### Gene accession numbers

Following gene accession numbers were used in the study: AT4G31805 (*AtPOLAR*), AT5G10890 (*At-POLAR-like1, AtPL1*), AT3G09730 (*AtPOLAR-like2, AtPL2*), AT5G61040 (*AtPOLAR-like3, AtPL3*); BdiBd21-3.3G0715200 (*BdPOLAR*), BdiBd21-3.1G0621800 (*BdPOLAR-like1, BdPL1*), BdiBd21-3.1G0412800 (*BdPOLAR-like2, BdPL2*), BdiBd21-3.3G0079200 (*BdPOLAR-like3, BdPL3*), BdiBd21-3.4G0050100 (*BdPOLAR-like4, BdPL4*), BdiBd21-3.3G0526300 (*BdPANGLOSS1, BdPAN1*), BdiBd21-3.5G0238000 (*BdYODA1, BdYDA1*), BdiBd21-3.5G0169200 (*BdBREVIS RADIX-solo, BdBRX-solo*); HORVU6Hr1G085100, HORVU7Hr1G030290, HORVU7Hr1G113890, HORVU6Hr1G028740, HORVU5Hr1G020410; Os02g55190, Os06g08520, Os06g43990, Os02g08470, Os12g38730, Os06g43240; Zm00001d052039, Zm00001d045105, Zm00001d015517, Zm00001d041525, Zm00001d046803; Potri.018G019400, Potri.006G263800, Potri.006G130600, Potri.015G057600; Solyc09T000143, Solyc03T002788.

#### Mature leaf phenotyping

For phenotyping of patterning defects in mature leaves, a mature third leaf of three-week-old soil-germinated plants was collected and fixed in a 7:1 solution of ethanol:acetic acid. Fixed and cleared leaves were rinsed in water, dried with a paper towel, and mounted on slides in Hoyer’s solution (15g gum arabic (Roth), 25mL milli-Q water, 200g chloral hydrate (Roth), 20mL glycerol (Thermo Fisher Scientific))^66^ for at least two days before imaging. The abaxial surface was then imaged using 20× and 40× objectives on a Leica DM2000 (Leica Microsystems) microscope with differential interference contrast (DIC).

Cell number of pavement cells, hair cells, subsidiary cells and stomatal complexes was counted manually using Fiji^62^, and the amount of clustered and disturbed cells was quantified. The phenotyping traits, such as hair and pavement cell clusters and proportion of affected subsidiary cells, were calculated by dividing the number of clustered or disturbed cells by the total cell number of each respective cell type. The cell index was calculated by dividing the count of each separate cell type or the count of stomatal complexes by the total cell count of the individual replica. For these measurements, the total number of five biological replicates was counted for wild type, *bdpl1-1*, and *bdpl1-2* alleles, and four biological replicates for the *bdpl1-3* allele. Total cell number was quantified for 4-5 fields of view per biological replica, stomatal complexes were counted as one cell.

For the cell density measurements, three fields of view were selected for three individuals of each genotype. For each field of view, the area was measured, and the total cell number was counted. The cell density was calculated by converting the μm^2^ into mm^2^ and dividing the total measured area by the total number of cells.

#### Generation of reporter constructs

Most constructs in this study were generated using the GreenGate modular cloning system^67^, which allows rapid assembly of six entry modules (pGGA-pGGF) into a destination vector through type IIS restriction-ligation reaction using BsaI-HFv2 and T4 DNA ligase. The destination vector pGGZ004^58^ was used for all final assemblies. The standard entry modules pGGA000, pGGB003, pGGC000, pGGD002, pGGE001, and pGGF000 were as described in^67^; pGGD_linker-mVenus was obtained from^60^, and pGGD009_linker-mCitrine was kindly provided by Jan Lohmann.

Genomic fragments, promoter regions, and coding sequences were amplified from *Brachypodium distachyon* Bd21-3 genomic DNA using Q5 High-Fidelity DNA Polymerase (New England Biolabs, NEB Cat.No. M0289S). PCR products were gel-purified (NucleoSpin Gel and PCR Clean-up Kit, Macherey-Nagel Cat.No. 740609.250) and cloned into BsaI-HFv2-digested (NEB, Cat. No R3733S) and dephosphorylated (Antarctic Phosphatase, NEB Cat. No. M0289S) GreenGate entry vectors (pGGA000-pGGF000). Ligation reactions were incubated overnight at 16°C with T4 DNA ligase (Thermo Fisher Scientific, Cat. No. EL0011). Positive clones were screened by colony PCR (OneTaq Polymerase, NEB Cat. No M0509L), verified by restriction digest and Sanger sequencing.

For the final reporter constructs 1.5 µL of each entry module (A-F) and 1.0 µL of the destination vector pGGZ004 were assembled in a 20 µL reaction containing 2.0 µL of 10× T4 ligase buffer, 0.5 µL T4 DNA ligase, and 0.5 µL BsaI-HFv2. Reactions were thermal-cycled 50 times between 37°C and 16°C (5 min each), followed by 5 min incubation at 50°C and 5 min at 80°C. To enhance ligation efficiency, 1 µL 10 mM ATP and 1 µL T4 ligase were added and incubated for 1 h at room temperature. 5µL of each reaction was transformed into chemically competent E. coli DH5α (NEB) cells, and positive clones were confirmed by Colony PCR, test digest and whole plasmid sequencing.

The BdPOLAR-LIKE 1 promoter (3,525 bp upstream region) was amplified using priDZ111/priDZ112 primers and cloned into pGGA000 to generate the pGGA_ BdPOLAR-LIKE1pro.

The BdPOLAR-LIKE 1 coding sequence (without stop codon) was amplified in two fragments to remove an internal BsaI site. The first fragment was amplified with priDZ113/priDZ144, and the second with priDZ143/ priDZ115. Both fragments were digested with BsaI-HFv2 and ligated with pGGC000 in a triple-ligation reaction to generate pGGC_BdPOLAR-LIKE1. Correct assembly was confirmed by colony PCR, restriction test, and sequencing.

For the cloning of myristoylation vector, priMR456 and priMR457 were annealed and phosphorylated with T4 Polynucleotide Kinase (NEB Cat.No. M0201L) to synthesise a small myristoylation peptide. pGGB000 was digested by BsaI and dephosphorylated. Finally, pGGB-myristoylation (MYR) was generated by ligating the phosphorylated and annealed primers into digested pGGB000. Construct was confirmed through analytical restriction digestion and Sanger sequencing.

Previously published entry modules pGGA_BdPO-LARpro, pGGA_ZmUbipro, pGGB003 (dummy), pGGD_ linker-mVenus, pGGE001, and pGGF_PvUbi2pro-HygR^33^ were used together with pGGA_BdPOLAR-LIKE1pro and pGGC_BdPOLAR-LIKE1 to generate promoter-swap constructs by introducing combinations of entry vectors into the destination vector pGGZ004 by GreenGate reaction.

For the BdBRX-solo translational reporter, three entry modules were generated: an 3.5Kb upstream promoter region, ending at the first codon, was amplified using priKJ16/priKJ17 and cloned into pGGA000 (pKJ08), the CDS (without stop codon) was amplified using priKJ18/ priKJ19 and cloned into pGGC000 (pKJ07), and the terminator (∼1 kb downstream region), was amplified with priKJ20/priKJ21 and cloned into pGGE000 (pKJ06). Final assembly was performed as described above, introducing the modules into the destination vector pGGZ004 to create *BdBRX-solop:BdBRX-solo-mCitrine*.

For the plasma membrane marker, the pGGC000_ PIP1;4 entry vector^61^, and the pGGD000_GSL-mScarlet (kindly provided by Karin Schumacher) were assembled with pGGA_ZmUbipro, pGGB003 (dummy) and pGGF_ PvUbi2pro-HygR to create ZmUbip:PIP1;4-GSL-mScarlet.

#### Generation of CRISPR/Cas9 constructs

CRISPR/Cas9 constructs were generated using two complementary systems: the high-efficiency GRF-GIF system^68^ and the multiplex Gateway-compatible CRISPR system^69^.

For single-gene targeting, CRISPR constructs were generated using the binary vector JD633_CRIS-PRhigheff^68^. The vector was digested with PaqCI (NEB, Cat.No R0745S) and dephosphorylated with Antarctic Phosphatase (NEB, Cat.No. M0289S). Geneious Prime (version 2025.2.2, GraphPad Software LLC d.b.a Geneious) was used to select candidate guide RNA (gRNA) with a high predicted activity score^70^. The generation of gRNAs was done by phosphorylating with T4 Polynucleotide Kinase (NEB, Cat.No. M0201L) and annealing of complementary oligonucleotides encoding gRNAs. Primer pairs priDZ91/priDZ135 and priDZ93/priDZ136 were used to synthesise gRNA1 and gRNA2, respectively. Annealed and phosphorylated oligonucleotides encoding gRNAs were ligated using T4 DNA ligase (Thermo Fisher Scientific, Cat.No. EL0011) with dephosphorylated and digested JD633_CRISPRhigheff vector, producing JD633_BdPO-LAR-LIKE1-gRNA1 and JD633_BdPOLAR-LIKE1-gRNA2 CRISPR/Cas9 constructs.

For dual-gene targeting, the pMDC32-based multiplex system^69^ was used. To generate pMDC32_Bd-POLAR-LIKE1-guide1/BdPOLAR-guide4, initially, Bd-POLAR-LIKE1-guide1 was synthesised by priDZ90 and priDZ91, BdPOLAR-guide4 was phosphorylated and annealed with priDZ88 and priDZ89, then both were ligated into linearised gRNA expression vector pYPQ131C and pYPQ132C separately. Next, pYPQ131C_BdPOLAR-LIKE1-guide1 and pYPQ132C_BdPOLAR-guide4, two gRNA expression cassettes, were cloned into pYPQ142 by Golden Gate reaction. At last, pYPQ142_Bd-POLAR-LIKE-guide1/BdPOLAR-guide4 was subsequently recombined with pYPQ167 into the destination vector pMDC32 through Gateway Assembly.

#### Generation of transgenic lines

Transformation and plant regeneration were performed as described in Zhang et al.^33^ In short, embryogenic calli were initiated from isolated embryos of Bd21-3, *bdpl1-1*, or *bdpolar-1* on callus induction medium (CIM: per L, 4.43 g Linsmaier & Skoog (LS; Duchefa), 30 g sucrose, 600 µL CuSO4 (1 mg/mL; Sigma/Merck), 500 µL 2,4-D (5 mg/mL in 1 M KOH; Sigma/Merck); pH 5.8; 2.3 g Phytagel (Sigma/Merck)). Media was refreshed by transferring calli to the new plates with fresh media and incubated in the dark at 28 °C for 1 week between transfers.

AGL1 *Agrobacterium tumenfaciens* carrying the desired construct was plated on the LB media with antibiotics for proper selection. Grown bacteria were scraped from selection plates and resuspended in liquid CIM, containing freshly added 2,4-D, 2.5 mg/mL; acetosyringone, 200 mM (Sigma/Merck); Synperonic PE/F68, 0.1% (Sigma/Merck). The transformation media was adjusted to an OD600 of 0.6. 50-100 calli were incubated in the transformation media for approximately 15 minutes at room temperature on a bench shaker (150rpm).

Transfected calli were partially dried out on sterile filter paper and co-cultivated for 3 days at room temperature in the dark. Calli were then transferred to the selection medium (CIM + Hygromycin, 40 mg/mL + Timentin, 200 mg/mL (Duchefa); Ticarcillin 2 Na / Clavulanate K) and incubated for 1 week at 28 °C in the dark. After 1 week, calli were moved to fresh selection plates and incubated 2 more weeks at 28 °C in the dark. Afterwards, calli were transferred to callus regeneration medium (CRM) (per L: 4.43 g Linsmaier & Skoog 30 g maltose (Sigma/Merck), 600 µL CuSO4 (1 mg/mL); pH 5.8; 2.3 g Phytagel; Timentin 200 mg/mL final; Hygromycin 40 mg/mL final; kinetin 0.2 mg/mL final (Sigma/Merck) and incubated at 28 °C, 16 h light / 8 h dark, PPFD 70-80 µmol m^-2^ s^-1^. Shoots > 1 cm typically form within 2-6 weeks.

Individual shoots were transferred to rooting cups containing rooting medium (per L: 4.3 g Murashige & Skoog + vitamins (Duchefa), 30 g sucrose; pH 5.8; 2.3 g Phytagel; Timentin 200 mg/mL final) and grown at 28 °C, 16 h light / 8 h dark, PPFD 70-80 µmol m^-2^ s^-1^ until roots formed. Rooted plants were transferred to soil and grown in a growth chamber under the above-mentioned growth conditions.

#### Crossing of Brachypodium plants

Crosses were performed following established procedures adapted for *Brachypodium distachyon* (Bd21-3 background). Plants were grown in soil for 4-6 weeks, depending on the growth rate of each line, until spikelets reached reproductive maturity. For pollen donor plants, 20 or more mature anthers were collected onto a microscope slide, embedded in a Petri dish with a wet paper towel and incubated at 28 °C for 10-50 min to induce dehiscence. Before crossing, the oldest unfertilised florets of recipient plants were selected and marked. Florets suitable for crossing displayed open and branched stigmas that were receptive to pollination. Selected florets were emasculated under a stereomicroscope by carefully removing all anthers using fine tweezers, then dehisced anthers were gently brushed across the receptive stigmas of emasculated florets. Pollinated florets were sealed with a small piece of medical tape over the lemma and palea to maintain closure until the full maturation of the seed.

*BdPL1p:BdPL1-mVenus* was crossed as the recipient with *ZmUBIp:PIP1;4-mScarlett* as the pollen donor, to generate *BdPL1p:BdPL1-mVenus;ZmUBIp:PIP1;4-mS-carlet* line used for the time-lapse experiment.

#### Measurement of cell area in developing epidermis

For quantitative analyses of cell size post-division, the second leaf of 5-6-day-old, WT, *bdpl1-1, bdpl1-1;BdPL1p:BdPL1-mCitrine* and *bdpl1-1;BdPL1p:*^*MYR*^*BdPL1-mCitrine* seedlings was used for imaging and stained with propidium iodide. For each genotype, five individual plants were analysed, with one representative field of view selected per leaf. The field of view was chosen within the developmental zone before the stage where SMC divisions had begun and all transverse ACDs were completed. 2D image segmentation and morphometric quantification were performed using MorphoGraphX^39^. Individual cell areas were extracted separately for each plant and combined across the five biological replicates per genotype to generate a cell size distribution.

#### Fluorescence intensity quantification and polarity analysis

The measurement of fluorescence intensity around a cell membrane was performed using a modified version of the POME pipeline^40^.

For each genotype, 3-5 biological replicates were analysed, with a total of 18-29 cells per line (Figures S4A-S4D and S5C-S5D). Isolated, individual cells were manually selected to avoid interference of the fluorescent signal from neighbouring cells.

The fluorescence intensity was measured along the cell membrane using a fixed reference angle of 0 radians, measured clockwise. The calculated angular coordinate ranged from 0 to 2π, covering the full 360° perimeter of each cell. For each calculated degree, the mean intensity of four consecutive pixels from the inner edge of the membrane outline was calculated.

For each cell, intensity values were normalised such that the maximum intensity was defined as 100% and the minimum as 0%. Normalised profiles were then aggregated per genotype, and the median intensity was calculated for each calculated angle across all cells of that line. Median intensity values were plotted against the angular coordinate to produce polarity domain profiles for each genotype. The standard deviation of raw values was plotted at the corresponding angle of colour-coded median intensity values as a line; coordinate axes for standard deviation lines were placed at 135° and 315°, with a scale from 0 to 100. The script for the adapted POME measurements and the generation of the polarity profile plots can be found here (https://github.com/raissig-lindner-lab/Korosteleva-et-al_2026_BdPL1).

The polarisation degree was quantified by calculating the ratio of fluorescence intensities at the apical (90°) and basal (270°) positions. Mean values of the four-pixel means at the closest calculated angles were used to compute the apical-to-basal polarity ratio for each cell.

#### Quantification and statistical analysis

Previously described measurements were statistically analysed with R (version 4.4.1) in RStudio (2025.05.0+496), and all plots were created with the ggplot2^71^ package. Other packages that were used for statistical analysis and plotting were: dplyr^72^, tidyverse^73^, scales, kableExtra, plotrix, quantreg, stargazer, emmeans, qPCR, ggpubr, rstatix and multicomp (for detailed information see www.r-project.org).

For statistical comparisons of short-range distribution (i.e. cell index, portion of clustered or disturbed cells, cell densities, fluorescence intensity ratios), we used one-way ANOVA, followed by Tukey’s HSD for multiple-comparison adjusted p-values. To show post-hoc groupings on the plots, compact letter displays (CLD) were generated from the Tukey table (multcomp/multcompView). For each genotype, the mean and standard deviation (SD) were calculated, and the data were visualised as boxplots with overlaid points (ggplot2) and CLD letters positioned above the boxes. Statistical significance was set at p < 0.05.

## Notes

### Competing Interest Statement

The authors have declared no competing interest.

