## Supplementary material for "A premitotic polarity program patterns the grass leaf epidermis": Figures S1-S7; Tables S1-S2

**Supplemental information**

**for**

**Korosteleva et al. (2026)**

This file contains:

Figures S1 - S7

Tables S1 - S2

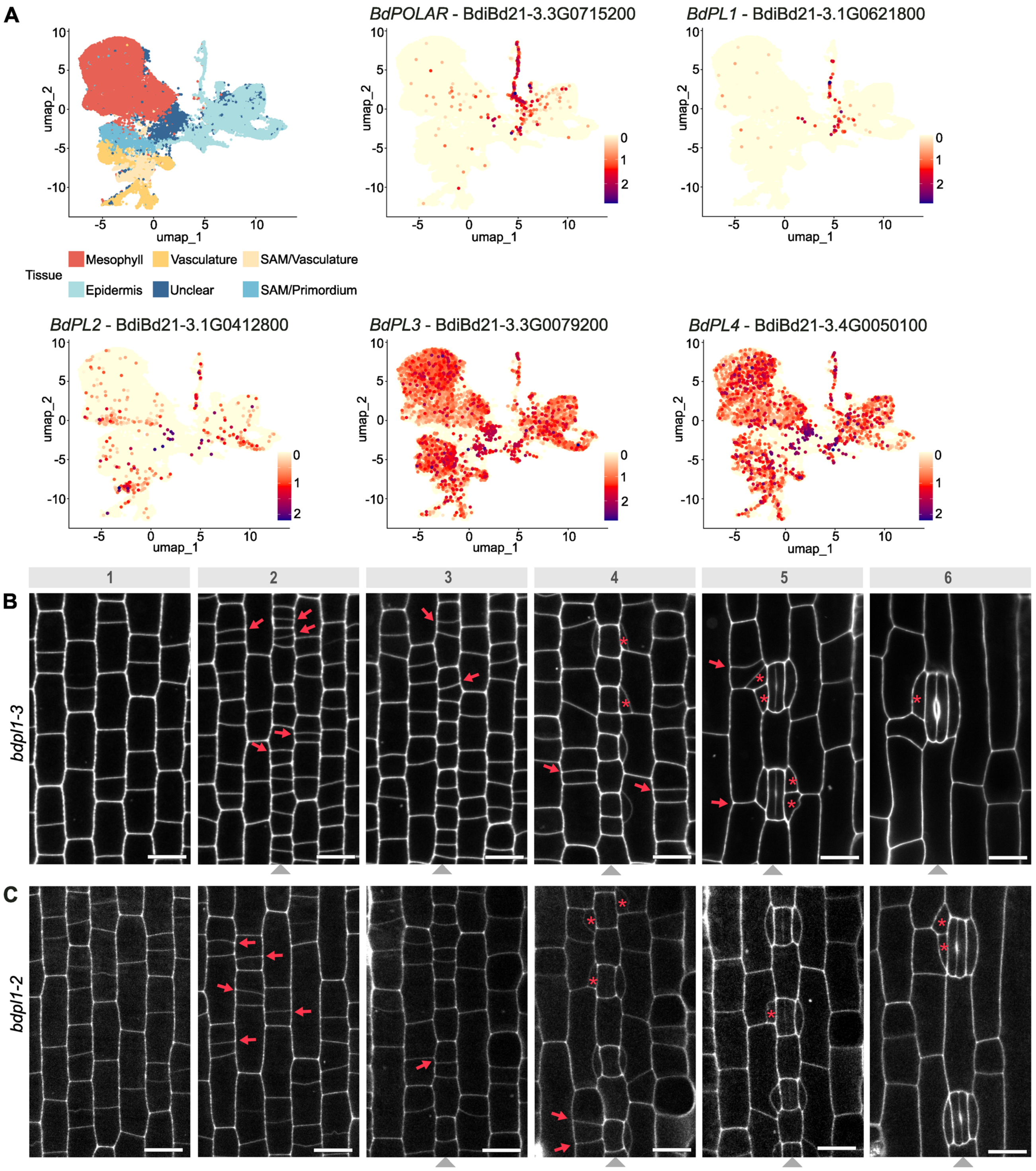


**Figure S1.** ***bdpl1* mutants show misoriented transverse asymmetric divisions**

**(A)** UMAP feature plots of the POLAR family genes in the young leaf of *Brachypodium dystachyon*[55]. Leaf tissue annotations for UMAP plots are indicated.

**(B-C)** Developmental zone of the 2$^{nd}$ leaf epidermis of 5-day-old seedlings of *bdpl1-3* (A) and *bdpl1-2* (B). Panels (1-6) show stages of epidermal development from undifferentiated protodermal cells (1) to mature stomatal complexes (6). Cell walls stained with propidium iodide (PI). Red arrows point at misoriented divisions, red asterisks mark disturbed subsidiary cells (SCs). Grey arrowheads indicate stomatal cell files. Scale bars, 10 µm.

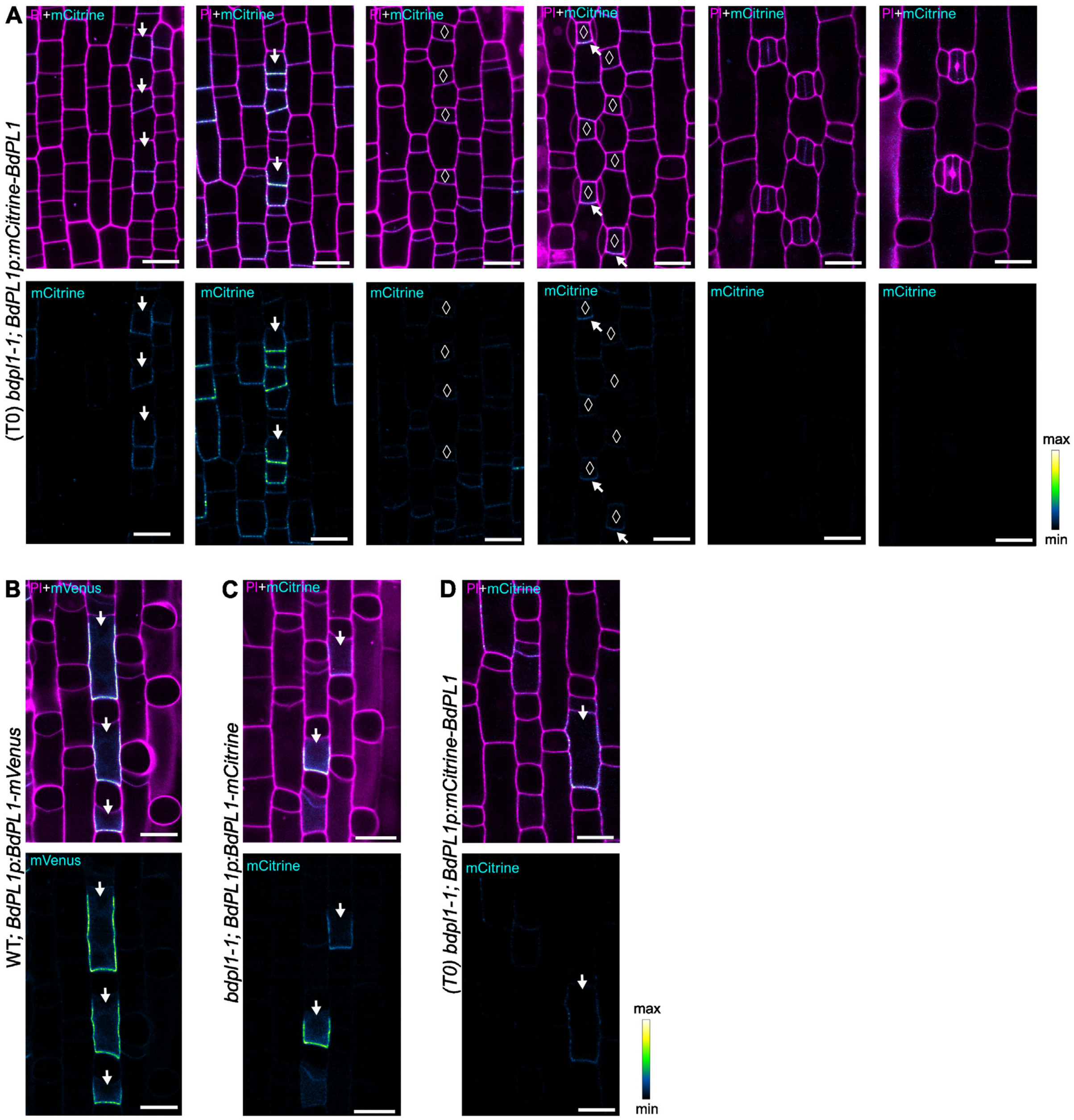


**Figure S2. Expression and subcellular localisation of different BdPL1 fusion proteins in wild-type and *bdpl1-1*.**
**(A)** Representative confocal images of *BdPL1p:mCitrine-BdPL1* (cyan) expression in *bdpl1-1* background in a leaf epidermis of T0 transgenic seedlings.
**(B-D)** Confocal images of BdPL1 fusion proteins showing pre-division polarisation in silica mother cells. *BdPL1p:BdPL1-mVenus* expressed in wild-type (WT) background (B), *BdPL1p:BdPL1-mCitrine* (C) and *BdPL1p:mCitrine-BdPL1* (T0, D) in *bdpl1-1* background. Upper panels show merged signals of mCitrine/mVenus (cyan) and PI-stained cell walls (magenta), lower panels show only mCitrine/mVenus signal ("Green Fire Blue" heatmap). White arrows indicate polarisation of BdPL1 fusion proteins, white diamonds indicate GMCs. Scale bars, 10 µm.

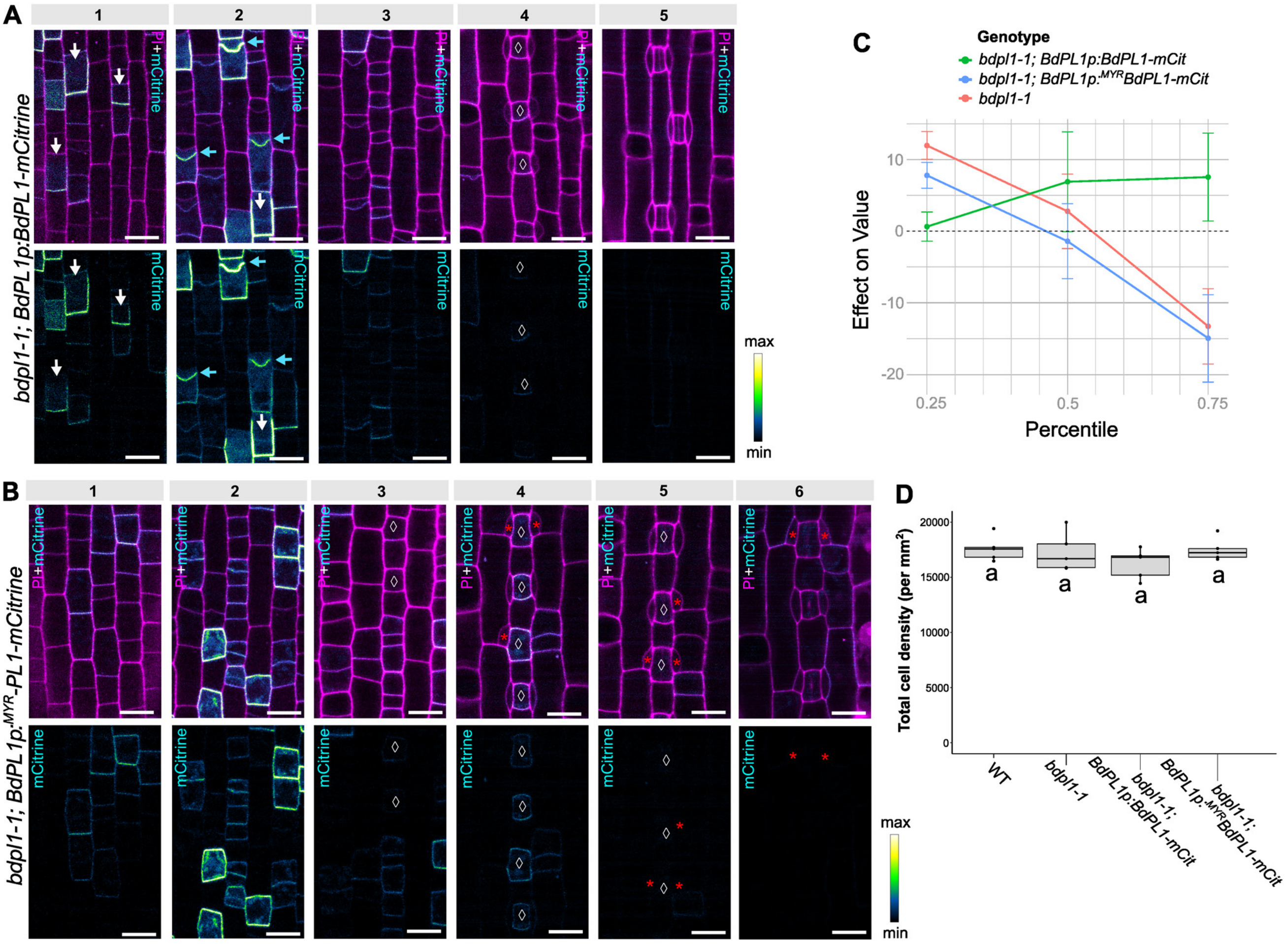


**Figure S3. Complementation lines and complementation analysis of *bdpl1-1*.**

**(A-B)** Developmental zone of the second leaf epidermis of 5-days-old seedlings expressing *BdPL1p:BdPL1-mCitrine* (A) and *BdPL1p:$^{MYR}$BdPL1-mCitrine* (B) in *bdpl1-1* background. Upper panels show merged signals of mCitrine (cyan) and PI-stained cell walls (magenta), lower panels show only mCitrine/mVenus signal ("Green Fire Blue" heatmap). White arrows indicate polarisation of BdPL1-mCitrine (A), cyan arrows point at relocalisation of BdPL1-mCitrine during transverse ACDs (A), white diamonds indicate GMCs, red asterisks indicate disturbed SCs (B). Scale bars, 10 µm.

**(C)** Effect of genotype on cell area distribution. Quantile analysis comparing cell area distributions at the 25$^{th}$, 50$^{th}$ and 75$^{th}$ quantiles between the intercept (wild type, WT) and *bdpl1-1*, *bdpl1-1;BdPL1p:BdPL1-mCitrine* and *bdpl1-1;BdPL1p:$^{MYR}$BdPL1-mCitrine*. Lines represent regression coefficients relative to WT; whiskers represent 95% confidence intervals

**(D)** Cell density in the second leaf epidermis at stage 2 (post-transverse ACDs), quantified as cells per mm². Genotypes as indicated. Boxes represent median and interquartile ranges; dots indicate biological replicates (n = 5 individuals per genotype). Letters below boxplots indicate statistical groupings determined by one-way ANOVA followed by Tukey's post hoc test (p-value < 0.05).

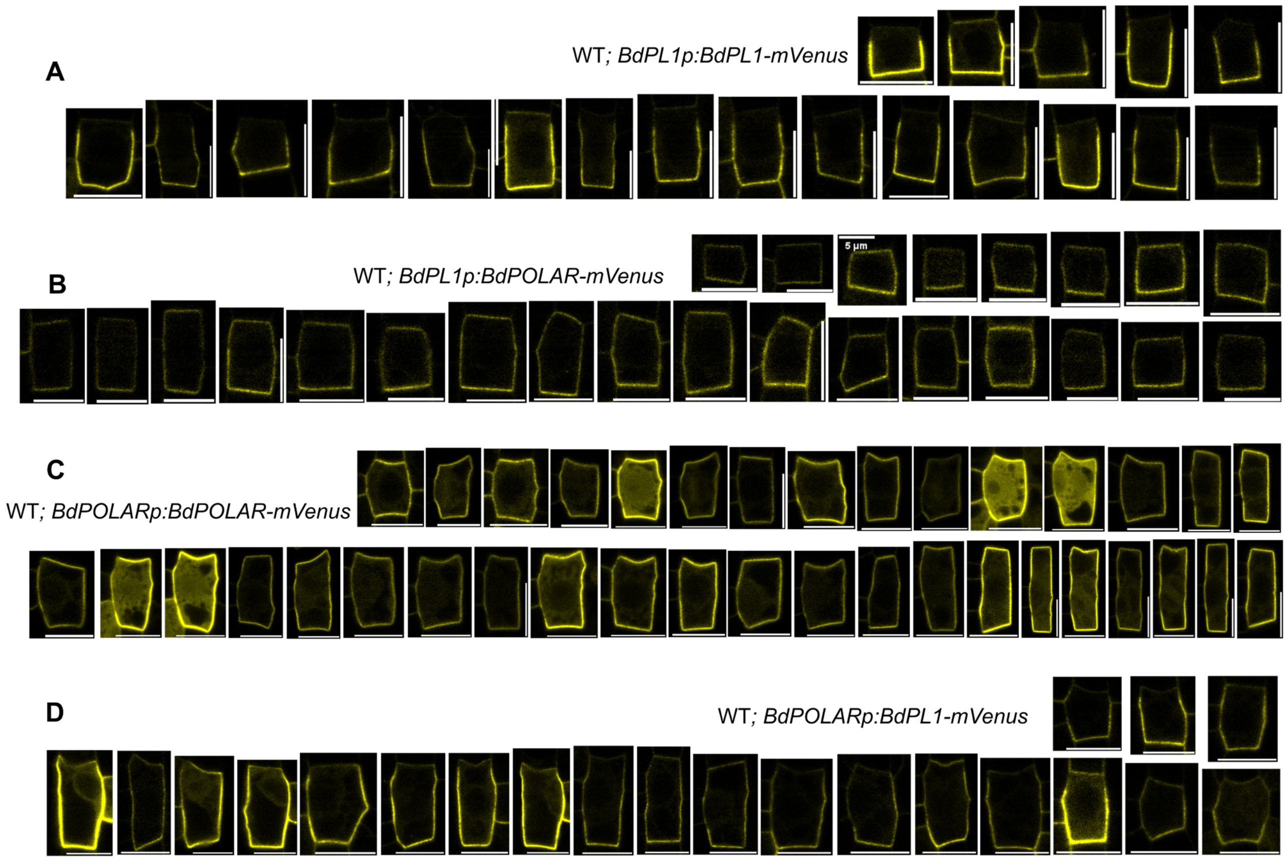


**Figure S4. Analysis of BdPL1 and BdPOLAR polarity domains in native and ectopic context.**
**(A-D)** Confocal single-plane images used for POME analysis of BdPL1-mVenus (yellow) and BdPOLAR-mVenus (yellow) under their native or reciprocal promoters in a wild-type (WT) background; *BdPL1p:BdPL1-mVenus (A), BdPL1p:BdPOLAR-mVenus* (B), *BdPOLARp:BdPOLAR-mVenus* (C), *BdPOLARp:BdPL1-mVenus* (D). Images are representative of ≥15 cells from three biological replicates. Scale bars, 10 and 5 (indicated) µm.

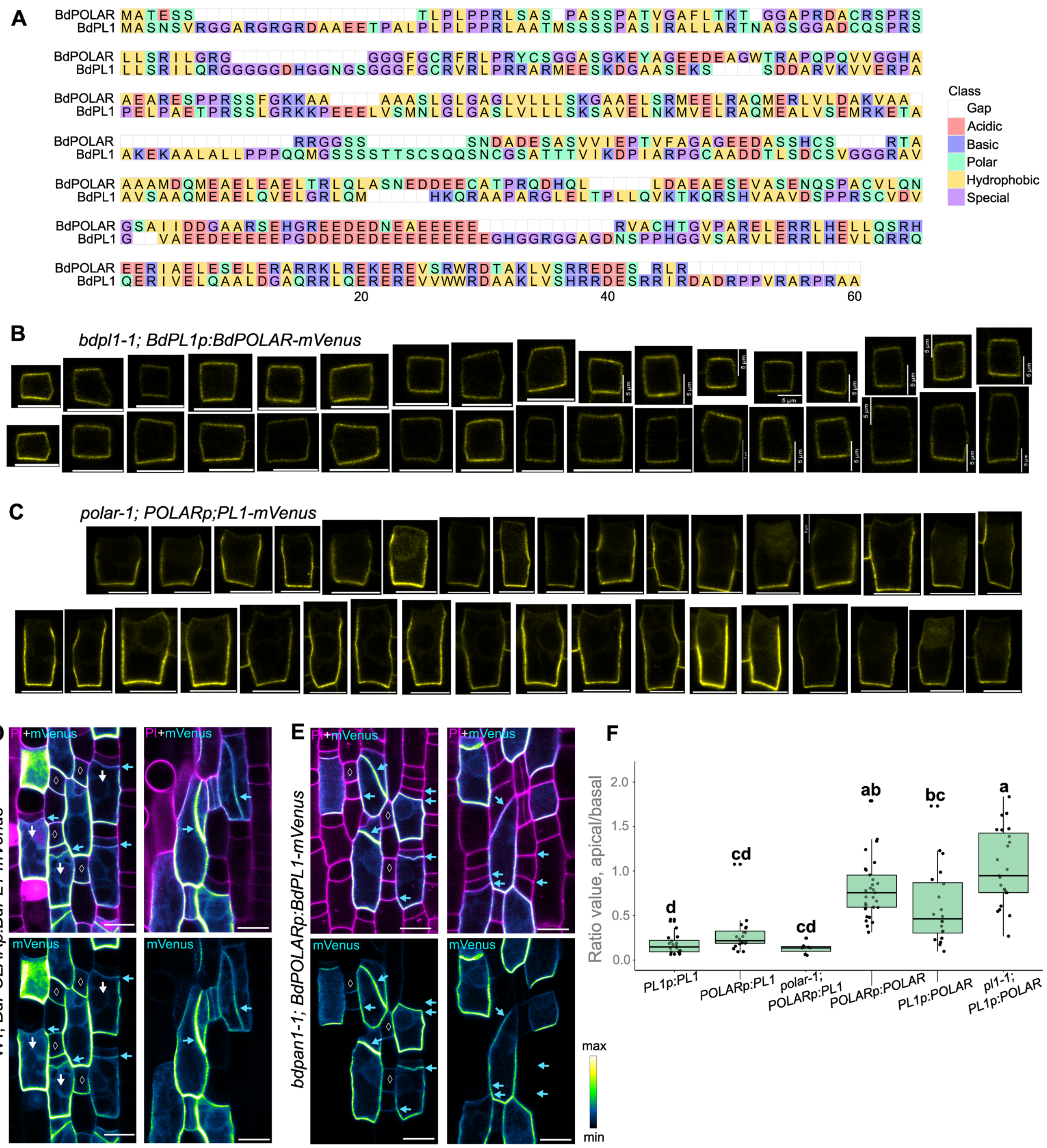

**Figure S5. BdPL1 and BdPOLAR share sequence similarity but display distinct polarity behaviours under reciprocal promoter control**

**(A)** Amino acid sequence alignment of BdPOLAR and BdPL1 proteins generated with MUSCLE [56] in MEGA12. Residues are colour-coded by biochemical properties according to the legend on the right.

**(B-C)** Confocal single-plane images showing the subcellular localisation of *BdPL1p:BdPOLAR-mVenus* in *bdpl1-1* background (B) and *BdPOLARp:BdPL1-mVenus* in *bdpolar-1* background (C). Reporter signals are shown in yellow; scale bars, 5 (indicated) and 10 µm.

**(D-E)** Confocal images of *BdPOLARp:BdPL1-mVenus* expression in wild-type (D) and *bdpan1-1* (E) backgrounds. Upper panels show merged signals for mVenus (cyan) and PI-stained cell walls (magenta), lower panels show only the mVenus signal ("Green Fire Blue" heatmap). White arrows indicate polarisation of BdPL1-mVenus, cyan arrows point at additional or misoriented divisions, white diamonds indicate guard mother cells (GMCs). Scale bars, 10 µm.

**(F)** Quantification of apical-basal fluorescence ratios for the indicated reporter lines. Grey dots indicate ratio values for individual cells, and boxes represent pooled values from 19-30 cells per genotype. Boxplots show the median and interquartile range, whiskers represent standard deviation (SD). Letters indicate statistically distinct groups based on one-way ANOVA followed by Tukey's post-hoc test (p-value < 0.05).

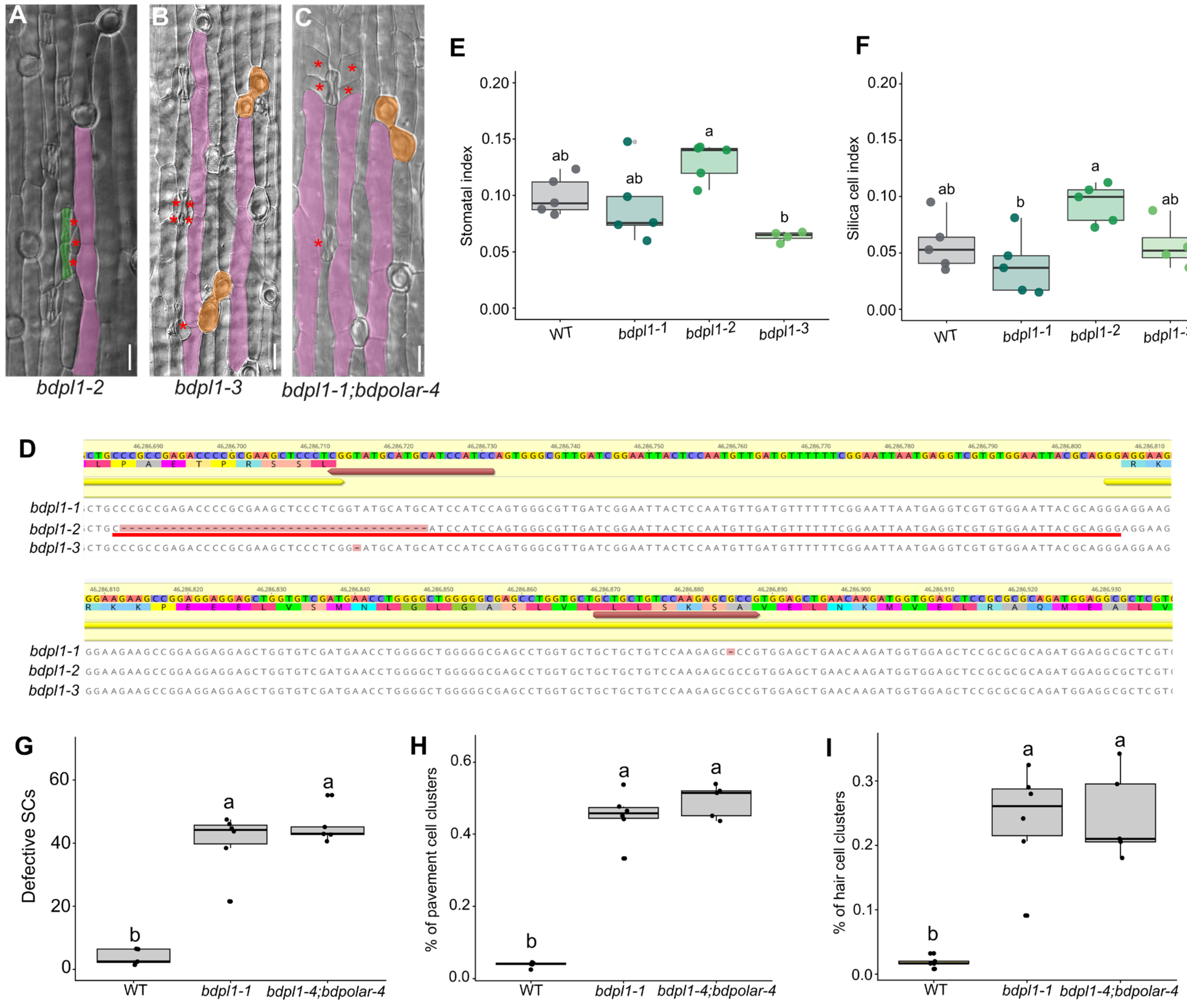


**Figure S6. Characterization of *bdpl1* alleles and the genetic interaction of *bdpl1* and *bdpolar.***

**(A-C)** Representative differential interference contrast (DIC) images of *bdpl1-2* (A), *bdpl1-3* (B), and *bdpl1-4;bdpolar-4* (C) leaf epidermis. Pavement cell clusters are false-coloured in pink, hair cell clusters are false-coloured in orange, guard cell clusters are false-coloured in green, and disturbed SCs are indicated with red asterisks. Scale bars, 20 µm.

**(D)** Sanger sequencing of three *bdpl1* alleles. *bdpl1-1* contains a single nucleotide deletion leading to a frameshift; *bdpl1-2* carries a 37 bp deletion causing a splice site removal and a 84bp in frame addition; *bdpl1-3* contains a 1 bp deletion outside of the first exon leading to a splice site mutation and a frameshift.

**(E)** Quantification of stomatal index (number of stomata complexes/(number of pavement cells + number of hair cells + number of silica cells + number of stomata complexes) in three *bdpl1* alleles and wild-type plants. Total cell number was quantified with 2-3 fields of view per individual for 5 individuals for WT, *bdpl1-1* and *bdpl1-2* and 4 individuals for *bdpl1-3*. Boxplots show the median and interquartile range, whiskers represent standard deviation (SD). Letters indicate statistically distinct groups based on one-way ANOVA followed by Tukey's post-hoc test (p-value < 0.05).

**(F)** Quantification of silica cell index (number of silica cells/(number of pavement cells + number of hair cells + number of silica cells + number of stomata complexes) in three *bdpl1* alleles and wild-type plants. Total cell number was quantified with 2-3 fields of view per individual for 5 individuals for WT, *bdpl1-1* and *bdpl1-2* and 4 individuals for *bdpl1-3*. Boxplots show the median and interquartile range, whiskers represent standard deviation (SD). Letters indicate statistically distinct groups based on one-way ANOVA followed by Tukey's post-hoc test (p-value < 0.05).

**(G-I)** Quantification of epidermal patterning defects in *bdpl1-1* and *bdpl1-4;bdpolar-4* mutants. Quantification of defective subsidiary cells (G), proportion of clustered pavement cells (H), and proportion of clustered hair cells (I). Boxes represent median and interquartile ranges; dots indicate biological replicates (n = 4-6 individuals per genotype). Letters indicate statistically distinct groups determined by one-way ANOVA followed by Tukey's post hoc test (p-value < 0.05).

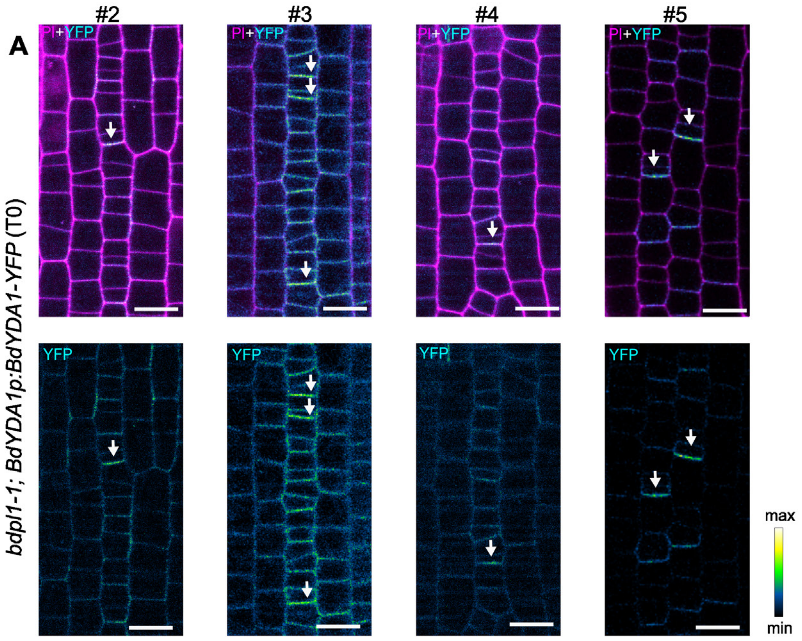


**Figure S7. Independent *BdYDA1-YFP* lines show consistent post-division polarisation in *bdpl1*.**

**(A)** Representative confocal images of leaf epidermis of four additional, independent T0 lines expressing *BdYDA1p:BdYDA1-YF*P in the *bdpl1-1* background. Upper panels show merged signals of YFP (cyan) and PI-stained cell walls (magenta), lower panels show only YFP signal (“Green Fire Blue” heatmap). White arrows indicate post-division polarisation of BdYDA-YFP. Scale bars, 10 µm.

**Table S1.** Deviation of estimated values and their statistical significance.

| Genotype | Estimate | Std. Error | P-value |
|---|---|---|---|
| Lower quartile (25th percentile) | | | |
| WT (Intercept) | 23.53050 | 0.56224 | 0.00000 |
| *bdpl1-1; BdPL1p:BdPL1-mCitrine* | +0.63030 | 1.04060 | 0.54474 |
| *bdpl1-1; BdPL1p:MYRBdPL1-mCitrine* | +7.79130 | 0.91912 | 0.00000 |
| *bdpl1-1* | +11.95600 | 0.99087 | 0.00000 |
| Median quartile (50th percentile) | | | |
| WT (Intercept) | 46.67670 | 2.38622 | 0.00000 |
| *bdpl1-1; BdPL1p:BdPL1-mCitrine* | +6.90000 | 3.54870 | 0.05192 |
| *bdpl1-1; BdPL1p:MYRBdPL1-mCitrine* | -1.40930 | 2.66768 | 0.59733 |
| *bdpl1-1* | +2.77020 | 2.65382 | 0.29662 |
| Upper quartile (75th percentile) | | | |
| WT (Intercept) | 84.44600 | 2.00684 | 0.00000 |
| *bdpl1-1; BdPL1p:BdPL1-mCitrine* | +7.54880 | 3.13699 | 0.01616 |
| *bdpl1-1; BdPL1p:MYRBdPL1-mCitrine* | -14.95690 | 3.10629 | 0.00000 |
| *bdpl1-1* | -13.28670 | 2.66889 | 0.00000 |

**Table S2.** Primers used in this study.

| Sequence | Name | Description |
|---|---|---|
| AACAGGTCTCAACCTTCACGTACGCTTAAGAAGGC | priDZ111 | FP:Amplifing BdPolar-like promoter |
| AACAGGTCTCATGTTCGATCCGATCCTTCAATTAC | priDZ112 | RP:Amplifing BdPolar-like promoter |
| AACAGGTCTCAGGAGCTGCCCGCCGAGTCCC | priDZ143 | FP:Amplifing BdPolar-like mutate BsaI |
| AACAGGTCTCACTCCGGCGCGGGCCTCTCGA | priDZ144 | RP:Amplifing BdPolar-like mutate BsaI |
| AACAGGTCTCAGGCTCCATGGCGAGTAATTCTGTT | priDZ113 | FP:Amplifing BdPolar-like CDS (TCC makes Ser) RP:priDZ144 |
| AACAGGTCTCACTGAAGCGGCGCGCGGCCGAGC | priDZ115 | RP:Amplifing BdPolar-like CDS (without stop codon) FP:priDZ143 |
| AACA ATGGGCAACAAATGTTGCAGCAAGCGACAGGA-TACCATGGCC | priMR456 | Myr-peptide |
| AGCCGGCCATGGTATCCTGTCGCTTGCTGCAACAT-TTGTTGCCCAT | priMR457 | Myr-peptide |
| TGGCACCCCTCTTTCATTTC | priDZ116 | Sequencing primer: BdPOLAR-like promoter FP |
| CCTGAGTTTATCACGGTCCG | priDZ117 | Sequencing primer: BdPOLAR-like promoter FP |
| AGGGCTAAACTTGTCCATGC | priDZ121 | Sequencing primer: BdPOLAR-like promoter RP |
| CTCTGGTGTTTTCTCCTCCG | priKJ14 | sequencing primer: POLAR-like1 promoter FP |
| CAGTTGGATTGCTGCGAAC | priKJ15 | sequencing primer: POLAR-like1 CDS RP |
| AACAGGTCTCAACCTCCCTTGTAGGCTCCGAGGGC | priKJ16 | Amplifying BRX-Solo promoter (BdiBd21-3.5G0169200) |
| AACAGGTCTCATGTTCTCGGCCCGGATCGCTCC | priKJ17 | Amplifying BRX-Solo promoter (BdiBd21-3.5G0169200) |
| AACAGGTCTCAGGCTCCATGCACGCGTGCTTCCAT | priKJ18 | Amplifying BRX-Solo CDS (BdiBd21-3.5G0169200) |
| AACAGGTCTCACTGAATTTATGTGCAAAGAACGACAC | priKJ19 | Amplifying BRX-Solo CDS (BdiBd21-3.5G0169200) without stop codon |
| AACAGGTCTCACTGCATAGAACTGATTTTTCTTTTCC | priKJ20 | Amplifying BRX-Solo terminator (BdiBd21-3.5G0169200) |
| AACAGGTCTCATAGTGTTCTGTTAATCCTGCAC | priKJ21 | Amplifying BRX-Solo terminator (BdiBd21-3.5G0169200) |
| CCCAAATCATCTATGGCGAG | priKJ22 | Sequencing BRX-Solo promoter |
| GCCACATACTTCTTAGGTTTCA | priKJ23 | Sequencing BRX-Solo promoter |
| GTTCTGCAATTTCGTCACC | priKJ24 | Sequencing BRX-Solo promoter |
| GCCATGTCGCTCAAGGTG | priKJ25 | Sequencing / cPCR primer BRX-Solo CDS |
| CGACACTTCCTTGGAGCAAC | priKJ26 | Sequencing / cPCR primer BRX-Solo terminator |
| CGAGGATGGAGGAGAGCAAG | priAK005 | FP for genotyping bdpl1 gRNA2 |
| GCGATGGGGTCCTTGATCAC | priAK006 | RP for genotyping bdpl1 gRNA2 |
| gtgtGCTGCTGTCCAAGAGCGCCG | priDZ90 | BdPOLAR-like-gR1-OsU6-F (guide 1) |
| aaacCGGCGCTCTTGGACAGCAGC | priDZ91 | BdPOLAR-like-gR1-OsU6-R (guide 1) |
| ACTTGGCTGCTGTCCAAGAGCGCCG | priDZ135 | BdPOLAR-like-gR1-OsU6-F(guide 1) used in GRF4–GIF1 system |
| aaacCGGTATGCATGCATCCATCC | priDZ93 | BdPOLAR-like-gR2-OsU6-R (guide 2) |
| ACTTGGGATGGATGCATGCATACCG | priDZ136 | BdPOLAR-like-gR2-OsU6-F(guide 2) used in GRF4–GIF1 system |
| gtgtGCTGCTATCCAAGGGCGCGG | priDZ88 | BdPOLAR-gR3-OsU6-F (guide 4) |
| aaacCCGCGCCCTTGGATAGCAGC | priDZ89 | BdPOLAR-gR3-OsU6-R (guide 4) |

**Table S3.** Cell count in mature leaf epidermis.

*see separate file "TableS3.xlsx"*
